# Plasmids encode and can mobilize onion pathogenicity in *Pantoea agglomerans*

**DOI:** 10.1101/2024.08.01.606178

**Authors:** Gi Yoon Shin, Jo Ann Asselin, Amy Smith, Brenna Aegerter, Teresa Coutinho, Mei Zhao, Bhabesh Dutta, Jennie Mazzone, Ram Neupane, Beth Gugino, Christy Hoepting, Manzeal Khanal, Subas Malla, Claudia Nischwitz, Jaspreet Sidhu, Antoinette Machado Burke, Jane Davey, Mark Uchanski, Michael L. Derie, Lindsey J. du Toit, Stephen Stresow, Jean M. Bonasera, Paul Stodghill, Brian Kvitko

**Affiliations:** Department of Plant Pathology, University of Georgia, Athens, GA, U.S.A.; Emerging Pests and Pathogens Research Unit, Robert W. Holley Center for Agriculture and Health, Agricultural Research Service, United State Department of Agriculture, Ithaca, NY, U.S.A.; University of California Cooperative Extension, Stockton, CA, U.S.A.; Department of Biochemistry, Genetics, and Microbiology, University of Pretoria, Pretoria, South Africa; Department of Plant Pathology, College of Plant Protection, China Agricultural University, Beijing, China; Department of Plant Pathology, University of Georgia, Tifton, GA, U.S.A.; Department of Plant Pathology and Environmental Microbiology, The Pennsylvania State University, University Park, PA, U.S.A.; Cornell Vegetable Program, Albion, NY, U.S.A.; Texas A&M AgriLife Research and Extension Center, Uvalde, TX, U.S.A.; Department of Biology, Utah State University, Logan, UT, U.S.A.; University of California Cooperative Extension, Bakersfield, CA, U.S.A.; Eagle Eye Produce, Iona, ID, U.S.A.; Department of Horticulture and Landscape Architecture, Colorado State University, Fort Collins, CO, U.S.A.; Department of Plant Pathology, Washington State University, Mount Vernon, WA, U.S.A.; Department of Horticulture, Michigan State University, Lansing, MI, U.S.A; Plant Pathology

## Abstract

*Pantoea agglomerans* is one of four *Pantoea* species for which strains have been reported in the United States to cause bacterial rot of onion bulbs. However, not all *P. agglomerans* strains are pathogenic to onion. We characterized onion-associated strains of *P. agglomerans* to elucidate the genetic and genomic signatures of onion-pathogenic *P. agglomerans*. We collected >300 *P. agglomerans* strains associated with symptomatic onion plants and bulbs from public culture collections, research laboratories, and a multi-year survey in 11 states in the USA. Genome assemblies were generated for 87 *P. agglomerans* strains that showed a range in onion virulence phenotypes. Combining the 87 genome assemblies with 100 high-quality, public *P. agglomerans* genome assemblies identified two well-represented and well-supported *P. agglomerans* phylogroups. Strains causing severe symptoms on onion leaves and bulbs were only identified in Phylogroup II and encoded the HiVir biosynthetic cluster for the phytotoxin pantaphos, supporting the role of HiVir as a crucial pathogenicity factor. Using a MASH-based plasmid classification system, the *P. agglomerans* HiVir cluster was determined to be encoded in two distinct plasmid contexts: 1) as an accessory gene cluster on a conserved *P. agglomerans* plasmid (pAggl), or 2) on a mosaic cluster of plasmids common among onion strains (pOnion). Analysis of closed genomes of *P. agglomerans* revealed that the pOnion plasmids harbored *alt* genes responsible for encoding tolerance to the thiosulfinate defensive chemistry in *Allium* spp. Additionally, many of these pOnion plasmids harbored *cop* gene clusters, which confer resistance to copper. However, the pOnion plasmids encoded the HiVir cluster less frequently. We demonstrated that the pOnion plasmid pCB1C, encoding HiVir and *alt* clusters as well as an intact conjugative type IV secretion system (T4SS), can act as a natively mobilizable pathogenicity plasmid that transforms *P. agglomerans* Phylogroup I strains, including environmental strains, into virulent pathogens of onion. This work indicates a central role for plasmids and plasmid ecology in mediating *P. agglomerans* interactions with onion plants, with potential implications for onion bacterial disease management.

## Introduction

*Pantoea agglomerans*, within family Erwiniaceae of the order Enterobacterales, is known for its widespread distribution and presence in diverse ecological contexts. Strains of *P. agglomerans* are ubiquitous in environments and frequently are isolated from animals, water, and soil. *P. agglomerans* strains also are common endophytes, epiphytes, and rhizosphere inhabitants of numerous plant species [Alkaabi et al. 2020; Andreote et al. 2008; Sulja et al. 2022; Walterson and Stavrinides 2015; Zhang et al. 2010]. In the phyllosphere, *P. agglomerans* strains may exhibit antagonistic effects against various plant pathogens and have been studied for biocontrol purposes [Costa et al. 2001; Giddens et al. 2002; Greiner and Winkelmann 1991]. Moreover, some strains can colonize the rhizosphere of rice (*Oryza sativa* L.) [Feng et al. 2006] and wheat (*Triticum aestivum* L.) [Soluch et al. 2021], promoting plant growth through nitrogen fixation and zinc solubilization, respectively. *P. agglomerans* strains are occasionally isolated as opportunistic wound pathogens in humans [Cheng et al. 2013; Cruz et al. 2007], including the type strain *P. agglomernas* LMG 1286^T^ (DSM 3493^T^=ATCC 27155^T^=NCTC 9381^T^) recovered from a human knee laceration. [Gavini et al. 1989].

*P. agglomerans* strains can also form pathogenic relationships with plants. *P. agglomerans* strains have been reported to cause disease in economically important monocot and dicot hosts, including onion (*Allium cepa* L.) [Hattingh and Walters 1981], rice [Lee et al. 2010; Xie 2001], Chinese cabbage (*Brassica rapa* L. subsp. *pekinensis*) [Guo et al. 2020], mango (*Mangifera indica* L.) [Gutiérrez-Barranquero et al. 2019] and gypsophila (*Gypsophila paniculata* L.) and beet (*Beta vulgaris* L.) [Burr et al. 1991; Cooksey 1986]. For gall-forming strains such as *P. agglomerans* pathovars *gypsophilae* 824–1 and *betae* 4188, pathogenicity is attributed to acquisition of the pPATH plasmid containing the genes for a *hrp1* type III secretion system (T3SS), host-specific effectors, and biosynthetic enzymes for plant hormones. The delivery of these effectors into plant cells leads to tumorigenic growth and gall formation [Barash and Manulis-Sasson 2007; Barash and Manulis-Sasson 2009; Nissan et al. 2018].

Recent reports in the United States have documented the association of *P. agglomerans* with onion diseases [Edens et al. 2006; Hu 2019; Khanal et al. 2023; Tho et al. 2015]. Strains of mutiple *Pantoea* species, namely *P. ananatis*, *P. allii*, *P. stewartii* subsp. *indologenes* pv. *cepacicola*, and *P. agglomerans* and have been reported to cause onion center rot [Brady et al. 2011; Carr et al. 2013; Gitaitis and Gay 1997; Hattingh and Walters 1981; Stumpf et al. 2018]. Onion center rot is characterized by leaf and scape necrosis and water-soaked, necrotic lesions in onion bulbs [Hattingh and Walters 1981; Edens et al. 2006]. Infections are typically initiated through wounds or thrips feeding [Hattingh and Walters 1981; Edens et al. 2006] [Gitaitis et al. 2003] [Dutta et al. 2014]. Foliar symptoms of center rot may be evident in the field. Bulb decay may develop in the field or during storage, transportation, and post-harvest handling if bacteria are present in the bulb or neck after harvest. [Vahling-Armstrong et al. 2016; Walcott et al. 2002]. Efforts to manage bacterial diseases of onions involve a multifaceted approach that integrates cultural, chemical, and biological control strategies [Dutta and Gitaitis 2020; Koirala et al. 2023]. Copper-containing pesticides commonly are applied to target various bacterial pathogens of onions, but copper tolerant strains of *P. agglomerans* have been in onion fields in the U.S.A. [Tho et al. 2019].

Previous work with onion-pathogenic *P. ananatis* strains identified key genetic determinants, namely the HiVir (<u>Hi</u>gh <u>Vir</u>ulence) and *alt* (<u>al</u>licin tolerance) gene clusters associated with onion pathogenicity and virulence [Agarwal et al. 2021; Asselin et al. 2018; Stice et al. 2018; Stice et al. 2021]. The HiVir gene cluster has been previously identified as a crucial factor for production of tissue necrosis in onion leaves and bulbs by *P. ananatis* and some strains of *P. stewartii* subsp. *indologenes* pv. *cepacicola* [Asselin et al. 2018; Zhao et al. 2023; Shin et al. 2023]. The chromosomal HiVir gene cluster in *P. ananatis* shows evidence of recent acquisition by horizontal gene transfer [Asselin et al. 2018] and encodes a series of biosynthetic enzymes responsible for producing the phosphonate phytotoxin pantaphos [Polidore et al. 2021], which induces necrosis in onion tissues. The plasmid-borne *alt* gene cluster has been identified as contributing tolerance towards thiosulfinates, plant antimicrobial compounds characteristically produced by *Allium* spp. as a consequence of cellular decompartmentalization following necrosis of onion cells [Stice et al. 2020]. Based on public draft genome assemblies, some *P. agglomerans* strains carry virulence genes clusters similar to those previously characterized in onion-virulent *Pantoea ananatis* strains [Stice et al. 2020; Stice et al. 2021; Shin et al. 2023].

Given that not all *P. agglomerans* strains isolated from onions exhibit pathogenicity to onion, we conducted an extensive examination of *P. agglomerans* strains associated with onions to elucidate the genetic and genomic characteristics of onion-pathogenic strains of *P. agglomerans*. In this work, over 300 *P. agglomerans* strains were collected from symptomatic onions from an extensive field survey spanning 11 states in the U.S.A. over two years, as well as from public bacterial culture collections and research laboratories. We generated closed and draft genome assemblies for 87 *P. agglomerans* strains isolated primarily from onions, and conducted phylogenetic and comparative genomic analysis of these new assemblies combined with 100 high-quality publicly available *P. agglomerans* genome assemblies. This work also includes the publication of 19 closed genomes of *P. agglomerans*, which permitted the detailed analysis of replicon gene content and variation. We identified two well-supported and well-represented *P. agglomerans* phylogroups. Notably, the strains causing severe symptoms on onion leaves and bulbs harbored a HiVir gene cluster and were exclusively found in Phylogroup II, highlighting both the crucial role of HiVir as a pathogenicity factor in *P. agglomerans* and its biased distribution. The *P. agglomerans* HiVir gene cluster was found to be plasmid-borne with approximately 76% average nucleotide identity with the chromosomally encoded *P. ananatis* HiVir cluster. The *P. agglomerans* HiVir cluster was found in two distinct plasmid contexts.

Within a sub-lineage of closely related strains isolated from across the U.S.A. and from South Africa, the HiVir cluster was encoded on a conserved *P. agglomerans* plasmid (pAggl). In other strains, HiVir was encoded on a mosaic cluster of plasmids prevalent among onion isolates (pOnion), which were further divided into three types. When present in *P. agglomerans* strains, the *alt* genes were found only on pOnion plasmids, and *cop* gene clusters encoding resistance to copper often were co-associated with *alt*. The pOnion plasmid pCB1C, encoding HiVir and *alt* clusters along with an intact conjugative type IV secretion system (T4SS), could act as a natively mobilizable pathogenicity plasmid, transforming *P. agglomerans* Phylogroup I strains, including environmental strains, into virulent onion pathogens. Thus, although no natively HiVir-encoding Phylogroup I strains were identified in our study, there do not appear to be any stringent barriers preventing this from occurring.

## Materials and Methods

### Isolation of bacterial strains from a national onion disease survey and preliminary identification using 16S rRNA gene sequencing

*P. agglomerans* strains conforming to the naming convention YYST####, indicating the two-digit year, two letter U.S.A. state abbreviation, and sequential isolate number (e.g., 20WA0189), were isolated as part of a United States national survey of bacteria associated with diseased onion plants and bulbs conducted during 2020 and 2021 in 11 onion-growing states (CA, CO, GA, ID, NM, NY, OR, PA, TX, UT, and WA). Isolates were recovered from symptomatic onion foliage and bulbs using standardized protocols. Briefly, small sections of leaf or bulb tissue (2 mm x 2 mm) were excised along the margin of the necrotic tissues using a sterile scalpel. The excised tissue was macerated in 100 µl of sterile water, and a sterile inoculation loop was used to streak macerate suspension onto nutrient agar (beef extract 3 g/L, peptone 5 g/L, agar 15 g/L, pH 7.0) or onion extract agar medium [Zaid et al. 2012]. After 48 hours of incubation at 28°C, pure cultures were generated through re-streaking and sub-culturing, as necessary. Pure cultures were preserved from nutrient broth (NB) cultures as 15% glycerol cryostocks stored at -80°C. Initial genus identification of isolates was conducted by 16S rRNA gene PCR assay and sequencing. DNA template for the PCR assay was prepared by combining 10 µl of overnight NB culture with 100 µl of sterile dH_2_O, followed by 95°C incubation for 10 minutes, and centrifugation of cell debris. See Supplemental Table S3 for 16S rRNA primer sequences and annealing temperatures. 16S rRNA gene amplicons sequences were determined by Sanger sequencing using either the forward or reverse PCR primer (Supplemental Table S3). Following sequencing and quality trimming, the bacterial strains were classified to genus using the SILVA Alignment, Classification and Tree service (https://www.arb-silva.de/aligner/). Strains were phenotyped further using the red onion scale necrosis (RSN) assay [Stice et al. 2018]. The study included 382 survey isolates representing a diversity of geographic locations in the U.S.A., sampling dates, onion tissue types, and RSN phenotypes, as well as 47 strains isolated from onion plants and bulbs and water sources near onion fields donated from the collection of Steven Beer at Cornell University, and 73 *Pantoea* strains previously isolated from onion or other sources by coauthors of this study. *Pantoea* strains (Supplemental Table S2) were screened using a species-specific PCR assay [Shin et al. 2022] (Supplemental Table S3) to identify *P. agglomerans* strains. PCR amplification and sequencing of the *infB* gene was used for secondary species confirmation for 32 *Panteoa* strains [Brady et al. 2008; Shin et al. 2022].

### Bacterial culturing

*P. agglomerans* and *Escherichia coli* strains were cultured in Luria-Bertani (LB) broth (containing 10 g/L tryptone, 5 g/L yeast extract, and 5 (University of Georgia, UGA) or 10 (U.S. Department of Agriculture, USDA) g/L NaCl) or agar (with 15 (UGA) or 16 (USDA) g/L agar) at temperatures of 28°C and 37°C, respectively. Antibiotics were added to the medium at the following final concentrations: gentamicin (Gm) at 10 µg/mL, kanamycin (Kan) at 50 µg/mL, streptomycin (Sm) at 100 µg/mL and rifampicin (Rp) at 50 µg/mL.

### Whole genome sequencing and assembly

Bacterial genomic DNA was extracted from overnight LB liquid cultures using a modification of the protocol from Miller et al. [1988] or either the Puregene Yeast/Bact. Kit B (Qiagen) or the Wizard HMW DNA Extraction Kit (Promega), following the manufacturer’s recommended Gram-negative bacterial DNA extraction protocol. For strain AR1a, DNA was extracted using a phenol: chloroform: isoamyl alcohol extraction and ethanol precipitation as in Asselin et al. [2021]. Genomic DNA was submitted to MicrobesNG, SeqCenter, or the Biotechnology Resource Center (BRC) Genomics Facility (RRID:SCR_021727) at the Cornell Institute of Biotechnology for Illumina short read sequencing. Oxford Nanopore long read sequencing was conducted either by Plasmidsaurus or in house at the USDA Agricultural Research Service (ARS) in Ithaca, New York. Genome sequencing at different facilities was conducted as described below.

#### 1. MicrobesNG

The samples were sequenced paired-end on an Illumina sequencer to generate 250 bp reads that were put through a standard analysis pipeline established at MicrobesNG (Birmingham, U.K). Briefly, the reads were mapped to the closest available reference genome identified by Kraken (https://github.com/DerrickWood/kraken), and assembled *de novo* using SPAdes (https://github.com/ablab/spades) to ensure quality assembly and 30X coverage. Resulting assemblies were annotated using Prokka (https://github.com/tseemann/prokka).

#### 2. SeqCenter (Previously MiGS)

The Illumina DNA Prep kit and IDT 10 bp UDI indices were used to prepare sample libraries that were sequenced on an Illumina NextSeq 2000. The 151-bp-long, paired-end reads were demultiplexed and subjected to quality control and adapter trimming with BCL Convert (v. 9.3) (Illumina).

#### 3. Long-read sequencing performed by Plasmidsaurus

The Oxford Nanopore Technologies Ligation Sequencing Kit version SQK-LSK114 was used to prepare samples for sequencing on GridION 10.4.1 flowcells (FLO-MIN114) with the "super accuracy" basecaller in MinKNOW.

#### 4. Short read sequencing performed by Cornell BRC Genomics Facility

For strains BH6c, SUH 1, and CB1, Illumina libraries were prepared using NEBNext Multiplex Oligos for Illumina (New England Biolabs), NEBNext Ultra FS DNA Library Prep Kit for Illumina (New England Biolabs), and AMPure XP beads (Beckman Coulter) according to manufacturer’s directions. Quality control of libraries was performed using BioAnalyzer (Cornell Biotechnology Resource Center Genomics Core) to calculate average library size and relative concentration of primers. Libraries were quantified using Qubit dsDNA HS assay (Thermo Fisher Scientific) and were pooled and submitted to Cornell BRC. For other strains sequenced at the BRC, purified genomic DNA was quantified using the Qubit dsDNA BR Assay and submitted. The BRC used the Nextera Flex kit (Illumina) to generate sequencing libraries that were sequenced using the MiSeq or NextSeq 500 instruments (Illumina) to generate ∼150-bp paired end reads.

#### 5. Long-read sequencing performed by Mt. Sinai Medical Center

AR1a was sequenced on a PacBio PSII (Pacific Bioscience) at the Icahn School of Medicine, Mount Sinai, using methods described in Mathers et al. [2015] and Asselin et al. [2021].

#### 6. Long-read sequencing performed by USDA-ARS

Genomic DNA quality was checked for potential degradation by agarose gel electrophoresis. DNA was quantified using a Qubit dsDNA BR Assay (ThermoFisher Scientific). Oxford Nanopore long-read DNA sequencing was conducted using the Spot-ON Flow Cell R9 version in a MinION Mk1B sequencing device. The library was prepared using the Rapid Barcoding Kit SQK-RBK and the flow cell was primed using the Flow Cell Priming Kit EXP-FLP002, according to the manufacturer’s instructions (Oxford Nanopore Technologies).

Basecalling was performed on the raw Nanopore data using Guppy (https://community.nanoporetech.com/downloads/guppy/). Indexed libraries were demultiplexed with “qcat” (https://github.com/nanoporetech/qcat). The specific version of each program used for each strain can be found with the NCBI SRA record for each sequence deposited.

### Genome Assembly and Annotation

#### Performed by Plasmidsaurus

Resulting raw reads were filtered with Filtlong (https://github.com/rrwick/Filtlong) to remove the lowest quality 5% of reads as well as reads with a length <2000 nucleotides. The filtered reads were subsampled to 400 Mb using rasusa (https://github.com/mbhall88/rasusa) and the bacterial genome assembly was carried out using the flye assembler (https://github.com/fenderglass/Flye) with -nano-hq and -read-error .02. Subsequent to assembly, polishing of the reconciled consensus was performed using medaka (https://github.com/nanoporetech/medaka). Finally, the genome annotations were made with bakta (https://github.com/oschwengers/bakta).

#### Performed at the University of Georgia (UGA)

FastQC (v. 0.11.8) [Andrews 2010] was used to evaluate the quality of the raw reads, and Trimmomatic (v. 0.39) [Bolger et al. 2014] was employed to filter out low-quality reads. The filtered paired-end reads were then assembled into contigs using SPAdes (v3.15.3) [Prjibelski et al. 2020], and the quality and completeness of the assemblies were assessed using QUAST (v. 5.0.2) [Gurevich et al. 2013] and BUSCO (v. 5.2.2) [Seppey et al. 2019].

#### Performed by the USDA-ARS in Ithaca, NY

The AR1a genome was assembled using Flye 2.6 [Kolmogorov et al. 2019] as described by Asselin et al. [2021].

The details of the assembly of each genome varied among strains. However, in general, each method followed the one suggested in the Trycycler documentation (https://github.com/rrwick/Trycycler): First, FiltLong (https://github.com/rrwick/Filtlong) was used to trim and filter the Nanopore reads. Next, filtered reads were subsampled randomly to 50x coverage (assuming a 5 Mb genome) to make 15 read subsets, and 5 draft assemblies each were generated using Flye [Kolmogorov et al. 2019], MiniPolish [Wick and Holt 2021], and Raven [Vaser and Šikić 2021] for a total of 15 draft assemblies. Trycycler [Wick 2020] was used to cluster and reconcile the 15 assemblies into a single consensus assembly. Medaka (https://github.com/nanoporetech/medaka) was used to polish the consensus assembly with the trimmed and filtered long reads. FASTP [Chen et al. 2018] was used to trim and filter the raw Illumina reads, and NextPolish [Hu et al. 2019] was used to polish the assembly with trimmed and filtered Illumina reads. Custom scripts were used to “normalize” the final assembly: (a) instances of the phi-X phage genome were removed; (b) the contigs were renamed, “chromosome”, “plasmidA”, “plasmidB”, etc., based on sequence length; (c) sequences were rotated to put, e.g., DnaA near position 1 on the positive strand.

The quality of each genome was assessed by annotating it with PGAP [Tatusova et al. 2016] and evaluating its proteome with BUSCO [Seppey et al. 2019]. Finally, an independent, “bottom-up” assembly was generated using Unicycler [Wick et al. 2017]. The Trycycler and Unicycler assemblies were compared using “dnadiff” from Mummer to identify potential assembly anomalies.

The specific version of each program used for each strain and can be found with the NCBI Assembly record for that genome.

### Data availability of genomes

The sequencing and assembly details, metadata, genome assemblies and corresponding annotations as well as the raw read data were deposited in the NCBI GenBank database under BioProject numbers PRJNA1069770 (UGA) and PRJNA642846 (USDA). The accessions of the deposited genomes can be found in Supplemental Table S6.

### Roary

In addition to 81 assembled genomes resulting from this study, 100 publicly available *P. agglomerans* genome assemblies were obtained from the NCBI GenBank database. The selection of these additional assemblies was based on their BUSCO scores and the availability of strain information, including the source, date, and location of isolation. The strain information for all 81 *P. agglomerans* genomes sequenced for this analysis can be found in Supplemental Table S6, and information for the 100 publicly available genomes used for this analysis can be found in Supplemental Table S7. Following the assembly, all 181 genomes were annotated for Gram negative bacteria using Prokka (v. 1.14.5) [Seemann 2014].

The core genes of the 181 annotated *P. agglomerans* genomes were identified with Roary (v. 3.13.0) [Page et al. 2015]. The concatenated nucleotide sequences of core genes were aligned with MAFFT [Katoh and Standley 2013] which was used to construct a phylogenetic tree using FastTree (v. 2.1.11) [Price et al. 2010]. The general time reversible (GTR) nucleotide substitution model was used to construct an approximately maximum likelihood phylogenetic tree, and the reliability of the internal branches was assessed by approximate likelihood-ratio test (aLRT) as the bootstrap method. The resulting tree was visualized and edited in iTOL (v. 6.8) [Letunic and Bork 2021] available at https://itol.embl.de/.

### Average nucleotide identity

FastANI (v. 1.3.3) [Jain et al. 2018] was employed to compute the pairwise average nucleotide identity (ANI) across the 187 *P. agglomerans* genomes. In addition to the Roary-analyzed 181 strains, six additional genomes of *P. agglomerans* from Steven Beer’s collection at Cornell University were included in the analysis. Following calculation, the ANI values were clustered hierarchically using the average linkage method and the one-minus Pearson correlation metric. Subsequently, a correlation plot was generated for all genomes using the ANI values by utilizing the Pearson correlation matrix option within Morpheus, a tool accessible at: https://software.broadinstitute.org/morpheus/.

### Phylogenetic analysis of a housekeeping gene, *gyrB*

Phylogenetic analysis of *gyrB* (DNA gyrase β-subunit) was conducted for 181 *P. agglomerans* strains by constructing a maximum-likelihood tree. The full-length nucleotide sequences of the *gyrB* gene (2,409 bp) were extracted from the 181 whole genome sequenced (WGS) *P. agglomeran*s strains using the BLAST function in Geneious Prime (v. 2013.1.2). The *gyrB* gene sequences from other *Pantoea* species were obtained from NCBI GenBank, with corresponding GenBank accession numbers provided in Supplemental Table S6. The *gyrB* gene sequences were aligned using the MAFFT alignment tool (v. 7.48), followed by trimming the alignment in Geneious Prime. The trimming process specifically targeted the regions corresponding to the partial *gyrB* sequence used for analysis by Bonasera et al. [2014] (745 bp) or Brady et al. [2008] (688 bp). The nucleotide substitution model for each alignment was chosen based on the Akaike Information Criterion (AIC) using the Smart Model Selection function in the online PhyML [Guindon et al. 2010] 3.0 tool. Branch support was evaluated through 1,000 replicate bootstrap analysis, and the resulting tree was visualized and edited in iTOL (v. 6.8). The partial *gyrB* sequence of *P. agglomerans* strains without genome sequences (non-WGS) was determined by PCR amplification and sequencing, as previously described [Bonasera et al. 2014]. See Supplemental Table S3 for primer sequences.

### Identification of characterized secondary metabolite and disease-associated gene clusters in sequenced genomes

Using BLAST+ [Camacho et al. 2009] (v. 2.11.0), the nucleotide sequences of the virulence and secondary metabolite gene clusters such as HiVir [Asselin et al. 2018], allicin tolerance (*alt*) gene cluster [Stice et al. 2018], copper tolerance gene cluster (this study), Pantocin A [Smits et al. 2019], *Pantoea* Natural Product 3 [Williams and Stavrinides 2020] and Herbicolin [Kamber et al. 2012] were searched against the genomes of 181 *P. agglomerans* strains. A cluster was considered present if 80% of the total gene cluster length was covered in the queried genome with a minimum of 70% nucleotide similarity. Following these criteria, we compiled a metadata table (Supplemental Table S9) indicating the presence and absence of these clusters. This table was then used as an input annotation file for iTOL (version 6.8, available at https://itol.embl.de/) to integrate it with the phylogenetic tree constructed using core genes.

### Selection of phylotype strains

To compute a set of potential representative strains for each phylogroup, we first computed the MASH [Ondov et al. 2016] distance for each pair of genomes in the phylograph using mashtree v. 1.4.6 [Katz et al. 2019]. Then, for each genome, we computed the average MASH distance to all other genomes within the phylogroup, generating a list of 10 strains. In a separate analysis, the gene group presence/absence matrix generated by the Roary analysis described above was used to compute the average Manhattan distance between each pair of closed genomes within each phylogroup. Within each phylogroup, strains with small MASH to other phylogroup strains, with large sets of shared orthologs, and that were also available in public culture collections were considered phylotype strains.

### Design and use of HiVir detection ‘PanHiVir’ primers

The full-length HiVir gene clusters (from *hvrA* to *hvrK)* from 31 HiVir-positive *P. agglomerans* strains with whole genome sequencing data, as well as from the genomes of other *Pantoea* species, such as *P. ananatis* 97-1R (GCA_002952035.2), LMG 2665^T^ (GCF_000710035.2), *P. allii* 20TX0020 (GCA_022585405.1), and *P. stewartii* subspecies *indologenes* pv. *cepacicola* PNA 03-3 (GCA_003201175.1), were downloaded from NCBI GenBank. The extraction and alignment were conducted using BLAST and MAFFT alignment functions in Geneious Prime (version 2013.1.2). Regions of perfect conservation within the HiVir gene cluster across these four *Pantoea* species were identified manually. Subsequently, we designed PanHiVir primers (Supplemental Table S3) targeting the *hvrD* gene to detect the HiVir gene cluster in these *Pantoea* species. These primers were used in colony PCR assays for a total of 502 *Pantoea* strains (Supplemental Table S2). The colony PCR reaction setup and protocol were similar to the *P. agglomerans*-specific PCR assay, with the exception of the annealing temperature (Supplemental Table S3).

### *copC* detection PCR assay and phylogenetic analysis

The nucleotide sequences of *copABCD* genes from a total of 65 *P. agglomerans* strains that contained copper tolerance gene clusters were aligned using MAFFT in Geneious Prime (v. 2013.1.2). For the design of *copC* detection primers, we selected conserved regions flanking the *copC* gene. Subsequently, we performed colony PCR assays using these *copC* primers to screen an additional 43 *P. agglomerans* strains that lacked whole genome sequencing. The details of the *copC* primer sequences and the annealing temperatures can be found in Supplemental Table S2. A total of 67 *copC* gene sequences (381 bp) from 65 *P. agglomerans* strains (comprising 53 with whole genome sequencing data and 12 without) were extracted and subjected to phylogenetic analysis following the procedure previously described for the *gyrB* gene.

### Clinker diagrams

The annotated HiVir and copper tolerance gene clusters from the closed genome strains of *P. agglomerans* were extracted and exported in GenBank format using Geneious Prime (v. 2013.1.2). The exported files were uploaded to the online comparative gene cluster analysis toolbox (available at: https://cagecat.bioinformatics.nl/tools/clinker) to generate a Clinker [Gilchrist and Chooi 2021] diagram. The minimum alignment sequence identity was set to a default value of 0.3 and the coloring scheme of diagram was changed manually.

### Analysis of plasmid sequences

The NCBI dataset program v. 15.23.0 (https://www.ncbi.nlm.nih.gov/datasets) was used to download all “complete” *P. agglomerans* (taxon = 549) genomes. In addition, the complete genomes assembled as part of this study were included to produce a single pool of complete genomes for analysis. To ensure that the genome annotations were consistent, each genome in the pool was reannotated with Prokka v. 1.14.6 [Seemann 2014]. In order to improve the quality of the output, each genome’s RefSeq or locally-generated PGAP [Tatusova et al. 2016] annotations, as appropriate, were provided to Prokka (--proteins GENOME.gbk) to use as a set of primary annotations. Next, BUSCO v. 5.5.0 [Seppey et al. 2019] was used to evaluate the quality of each genome and its Prokka annotation using the enterobacterales lineage database. Any genome with a “completeness” (C) score of < 95% or a “duplicated single-copy” (D) score of > 5% was removed from the pool. In order to ensure the genomes in the pool were identified correctly as *P. agglomerans*, a pair-wise ANI analysis of all genomes in the pool was performed using FastANI v. 1.33 [Jain et al. 2018]. Any genome that did not have an ANI score ≥ 95%, with respect to the *P. agglomerans* type strain (GCF_019048385.1), was removed from the pool.

Having constructed a pool of 35 high-quality, complete, consistently-reannotated *P. agglomerans* genomes (Supplemental Table S14), each genome’s sequence (.fna), proteome (.faa), and annotation (.gff) file was split using custom scripts into separate files for the chromosome and each individual plasmid. The MASH [Ondov et al. 2016] distance between each pair of replicons was computed using Mashtree v. 1.4.5 [Katz et al. 2019]. In addition to producing a table of the MASH distance between each part, Mashtree also produces a tree in Newick format. The mashtree_bootstrap.pl script with 100 reps was used to assign bootstrap values to the branches of the tree.

Using the MASH distances, replicons were assigned to plasmid clusters using the following method: a graph was constructed in which each replicon was represented by a node, and edges were created between pairs of nodes when the MASH distance was less than or equal to an arbitrary cutoff. For this analysis, we used a MASH distance of 0.05, which very loosely corresponds to a 95% identity. A custom script was used to compute connected components within this graph; each connected component represents a cluster of plasmids with similar *k*-mer statistics. Hereafter, we refer to these plasmid clusters as “plasters” and this analysis as the “plaster analysis”.

Some circular plasmid figures were generated using BRIG (BLAST Ring Image Generator) [Alikhan et al. 2011].

### Prediction of conjugative and mobilizable plasmids

For each of the 35 closed, high-quality *P. agglomerans* genomes, we used a custom script (gbk2gembase) to convert the annotation generated by Prokka into a proteome data file specifically formatted for MacSyFinder. Next, for each genome, we used this proteome data file as input to MacSyFinder v. 2.1.2 [Abby et al. 2014; Neron et al. 2022]. We analyzed each proteome with the CONJScan/Plasmids v. 2.0.1 models [Abby et al. 2016]. We specified that all submodels should be considered. We also specified that all replicons were circular.

### Panreplicon analysis

In order to determine whether or not the replicons in each plaster share a set of common, or core, genes, we ran Roary v. 3.13.0 [Page et al. 2015] with the set of individual replicons (not genomes) as input. In order to produce results for every gene group across all replicons, regardless of cardinality, Roary was run with the parameter, -cd 0.0. A custom script was used to post-process the Roary output in order to compute the percentage inclusion of each gene group within each plaster.

In the full-genome context, genes in the pangenome typically are classified based on the percentage of strains found to have a copy. For instance, the *hard core* of the pangenome may be defined as the set of genes found in 99 - 100% of the analyzed genomes, the *soft_core* as genes found in 95% or more of the genomes, the *shell* as the genes found in 15% or more of the genomes, and the rest of the genes as the *cloud*. To understand the gene complement of each plaster group’s replicons, we extended this definition as follows: the *hard core* of a plaster group is a set of genes found in 99 - 100% of the replicons that are members of this plaster group; similarly, the *soft core* of a plaster group are the genes found in 95% of the members of the plaster group; the definitions of *shell* and *cloud* for plaster groups are similar.

### Genetic manipulation

Spontaneous rifampicin resistant isolates were recovered by plating approximately 10^8^ CFU (200-300 µl of overnight shaken cultures in LB broth or a suspension in sterile water of cells recovered from LB plate cultures) onto LB rifampicin (LB Rp) amended agar medium. Antibiotic resistance was confirmed by dilution spotting suspensions onto both LB and LB Rp media and counting colonies to confirm similar growth rates. The rifampicin resistant strains also were assessed for RSN phenotypes compared to the wildtype (WT) strains prior to use in further experiments.

The in-frame deletion of the *hvrA* (phosphoenopyruvate mutase) gene from *P. agglomerans* AR1a was conducted following the established allelic exchange protocols previously utilized for *P. ananatis* [Stice et al. 2020; Shin et al. 2022] to exchange the *hvrA* gene with a synthesized DNA fragment (Supplemental Table S3) corresponding to the *hvrA* deletion cloned into pR6KT2G using BP clonase II. The Δ*hvrA* mutant was recovered in a two-step process by selection of GmR pR6KT2G single-crossover exconjugants followed by sucrose counter-selection to recover double-crossover mutants. The *P. agglomerans hvrA* deletion mutant strains were genotyped using PCR assays and verified by PCR amplicon sequencing (Supplemental Table S3).

Two KanR marked derivatives of the pCB1C plasmid in *P. agglomerans* CB1 were generated by allelic exchange using the pKNG101 vector [Kaniga et al. 1991] to insert the kanamycin resistance gene from pBS44 [Swingle et al. 2008] either between LD072_23365 and LD072_23370 (pCB1C-Akan) or replacing pCB1C LD072_23445 and LD072_23450 (pCB1C-BKan). DNA fragments were amplified by PCR assays and then joined and cloned into pBC SK^-^ using NEBuilder HiFi DNA Assembly (Supplemental Table S3). After sequencing confirmation, the correctly assembled fragments were sub-cloned into pKNG101 by SalI-HF and SpeI-HF restriction digest and T4 ligation. The KanR double cross-in mutants of CB1 pCB1C were recovered in a two-step process using selection of KanR SmR pKNG101 *P. agglomerans* CB1 single-crossover exconjugants, followed by sub-culturing without streptomycin selection, and screening for KanR SmS putative double-crossover mutants and PCR genotyping (Supplemental Table S3). Successful creation of *P. agglomerans* CB1 pCB1C-AKan and CB1 pCB1C-BKan was confirmed by Illumina sequencing at the Cornell Biotechnology Resource Center, as described above.

### Native mobilization of the Kan marked pCB1C virulence plasmid

*P. agglomerans* CB1 pCB1C-AKan or CB1 pCB1C-BKan donor strains and Rp-resistant AR8b, MMD61212-C, FC61912-B, and J22c recipient strains were suspended from fresh LB plate cultures to an OD_600_ of 0.4 in sterile water. Approximately 10^8^ CFU of each donor and recipient strain were combined, and the bacterial suspensions were then pelleted by centrifugation and re-suspended in 1 mL of LB. After 19 hours of incubation with shaking at 28°C, 0.2 mL of culture was plated onto LB Kan Rp agar medium to recover potential pCB1C-AKan or pCB1C-BKan exconjugants. Oxford Nanopore sequencing was conducted as described above to confirm the presence of pC-AKan in *P. agglomerans* AR8b, MMD61212-C, and FC61912-B exconjugants.

To estimate pC-AKan conjugation efficiency, water suspensions of the *P. agglomerans* CB1 pCB1C-Akan donor and the Rp-resistant MMD61212-C were prepared as described above and combined. The population of each parent in the combined suspension was determined by dilution series spot plating onto LB Rp and LB Kan agar medium. A 0.2 mL aliquot of the combined suspension was spread onto water agar medium and incubated overnight at 28°C.

Bacteria were recovered from the water agar in 0.4 mL of sterile water. The population of each parent after incubation on water agar was estimated by dilution series spot plating onto LB Rp and LB Kan agar media and 0.3 mL was plated onto LB Kan Rp agar medium to recover exconjugants. Putative exconjugants were screened by PCR assay to confirm the presence of pCB1C-AKan (Supplemental Table S3).

### Bacterial inoculum preparation

*P. agglomerans* strains selected for phenotyping assays were streaked onto LB or LM agar media (consisting of 10 g/L tryptone, 6 g/L yeast extract, 1.193 g/L KH_2_PO_4_, 0.6 g/L NaCl, 0.4 g/L MgSO_4·_7H_2_0, and agar 15 g/L) containing appropriate antibiotics from -80°C cultures and incubated overnight at 28°C. To create bacterial inoculum, a single colony of each strain was inoculated into 3 mL of LB or LM broth and incubated overnight with agitation at 200 rpm. Following incubation, the bacterial cells were harvested by centrifugation and the cell pellets were resuspended and adjusted to an OD_600_ of 0.2-0.3 using sterile 1X PBS buffer. Alternatively, bacteria were pulled from LB agar plates grown overnight at 28°C using sterile cotton-tipped applicators, and inoculated into autoclaved, high-purity water and adjusted to an OD_600_ of 0.2 - 0.3.

### Disease assays

Red onions were procured through the UGA dining services or purchased from local grocery stores in Athens, GA or Ithaca, NY. RSN assays were conducted for all *Pantoea* strains using a previously described protocol [Shin et al. 2023] with slight modifications. The onion scales were stab inoculated using either a sterile pipette tip or a toothpick that had been dipped in bacterial inoculum. Onion plants (cv. Century) were grown in the greenhouse in Athens, GA as described by Shin et al. [2023] and inoculated according to the cut tip inoculation method for foliar necrosis assays [Shin et al. 2023].

Red onion bulb necrosis assays were conducted using a modification of the onion bulb necrosis assay procedures described in Asselin et al. [2018]. Red onions without any visible wounds or external symptoms of infection were selected for the assay. After removing the dried outer scale layers, the surface of the onion bulb was disinfected by wiping with 70% ethanol. A sterilized, full-length, dissecting needle was used to puncture the shoulder on one side of the onion bulb, approximately 45 mm towards the center of the bulb. Following this, 10 µl of bacterial inoculum was pipetted into the puncture wound. Sterile 1X PBS buffer was used as a negative control treatment. Three separate onion bulbs were used per treatment. Bulbs were incubated upright at 28°C for one week, bisected vertically through the site of inoculation, and the cut surfaces was photographed.

For gain-of-function foliar disease assays with ex-conjugant strains, onion plants were grown as described by Bonasera et al. [2017]. Plants were moved to a growth chamber set at 28°C with 14 hours of light/day and ambient humidity, one day prior to inoculation. Bacterial inoculum was prepared in water to an OD_600_ = 0.2, as previously described. Plants were brought into the lab, and four plants were inoculated per treatment (strains or sterile water), and five leaves were inoculated per plant, starting with the youngest leaf > 10 cm above the soil line that was wide enough to be inoculated with a toothpick without detaching the tissue above the inoculation site. Leaves were inoculated approximately 10 cm above the soil line by stabbing the leaves with toothpicks dipped in inoculum. After inoculation, plants were returned to the growth chamber. Lesions on all leaves were measured 2, 3, 4, and 5 days after inoculation. The experiment was completed twice. Lesions from the fifth oldest inoculated leaves were not included in the analysis, as these leaves began to senesce during the experiment, and the lesions could not be distinguished from the naturally senescent tissue.

A linear mixed effects model was used to compare lesion size on day 5. The model included a fixed effect of treatment and a random effect of plant due to the fact that measurements were taken on 4 leaves per plant. Post-hoc comparisons among treatments were performed using Tukey’s honestly significant difference (HSD) test. A p-value < 0.05 was considered statistically significant.

### Copper tolerance assay

A copper tolerance assay was performed using 123 strains of *P. agglomerans* (81 whole genome sequenced (WGS) and 42 non-WGS strains) (Supplemental Table S25), following established protocols [Sundin 1989; Tho et al. 2019] with minor adjustments. Casitone Yeast Extract (CYE) agar medium, (composed of 1.7 g/L casitone, 0.35 g/L yeast extract, and 15 g/L agar) was supplemented with filter-sterilized copper sulfate (CuSO_4_·5H_2_O) at final concentrations of 50, 100, 150, or 200 µg/ml. To prevent formation of highly mucoid colonies that can interfere with interpretation of confluent growth, glycerol was omitted from the medium. As a control treatment, CYE medium without copper sulfate was used. The bacterial inoculum was prepared as previously outlined and arrayed in 96-well microtiter plates, and 1 µl of the inoculum was spotted onto CYE agar medium in gridded square petri plates using a multichannel pipettor. Afterward, the plates were dried in a biosafety cabinet and incubated at 28°C for 2 days. Bacterial growth at each copper concentration was observed and scored for confluent growth, non-confluent growth, or no growth. The assay was run with two technical replicates and was carried out three times.

## Results

### Strains and genomes used in this study

Six hundred and eighteen (618) *P. agglomerans* strains used throughout this study. Supplemental Table S1 lists the strains and analyses in which they were used.

To delineate the genetic and genomic relationships among *P. agglomerans* onion and other strains, whole genome Illumina short-read sequencing was conducted for 81 strains: 79 isolates from onions, one isolate from Carolina geranium (*Geranium carolinianum* L., a common weed in onion fields), and one isolate from eucalyptus (*Eucalyptus* L.). The set of selected *P. agglomerans* onion isolates had geographically diverse origins (77 strains isolated from 11 states in the USA, as well as 3 from South Africa, and 1 from Uruguay) and included onion pathogenic and non-pathogenic strains.

The QUAST and BUSCO assessment results of the assembled genomes can be found in Supplemental Table S4 and S5.

To characterize the plasmids of *P. agglomerans* and to understand better the relationship between plasmids and *Pantoea* pathology, 19 genome assemblies were closed via hybrid assembly of Illumina short reads and either Oxford Nanopore or PacBio long reads.

*P. agglomerans* is comprised of at least two phylogroups. Based on ANI analysis of the 81 new genome assemblies of *P. agglomerans* generated in this study combined with 100 public *P. agglomerans* genome assemblies selected based on genome completeness and metadata availability (location, source, and year of isolation) and the DSM 3494^T^ type strain, a clinical isolate recovered from a human knee laceration, all 181 genomes had above the 95% similarity threshold supporting their assignment as *P. agglomerans*.

Genomes annotated with Prokka resulted in a total of 27,204 annotated genes. Among these total observed genes, the Roary pangenome analysis identified 3,111 core genes present in 99% of the strains, along with 337 soft core genes found in 95% of the strains. Furthermore, the analysis revealed 1,652 shell genes present in 15 to 95% of the strains, and 22,104 cloud genes present in 1 to 15% of the strains.

The *P. agglomerans* core genome phylogeny supported the delineation of two well-represented phylogroups that included 175 of the 181 analyzed strains (Fig. 1). The presence of at least two well-supported phylogenetic divisions in *P. agglomerans* has been noted previously [Tambong 2019; Sulja et al. 2022; Crosby et al. 2023; Soluch et al. 2021]. Of the 181 sequenced *P. agglomerans* strains, six fell outside of these two well-represented phylogroups. We designated the phylogroup containing the *P. agglomerans* DSM 3494^T^ Type strain clinical isolate as Phylogroup I, and the second well-represented phylogroup as Phylogroup II. Although *P. agglomerans* DSM 3494^T^ is supported as a member of Phylogroup I, it is distantly related to other Phylogroup I strains based on this analysis Based upon average MASH distance and the size of shared ortholog sets between all pairs of strains within each phylogroup and availability of strains in public culture collections, we suggest *P. agglomerans* CFBP8784, isolated in France from radish (*Raphanus sativus* L.) seed, and AR5 (NRRL #B-65700), isolated from a diseased onion bulb grown in the state of Oregon U.S.A., as the phylotype strains for Phylogroups I and II, respectively.

**Figure 1.**
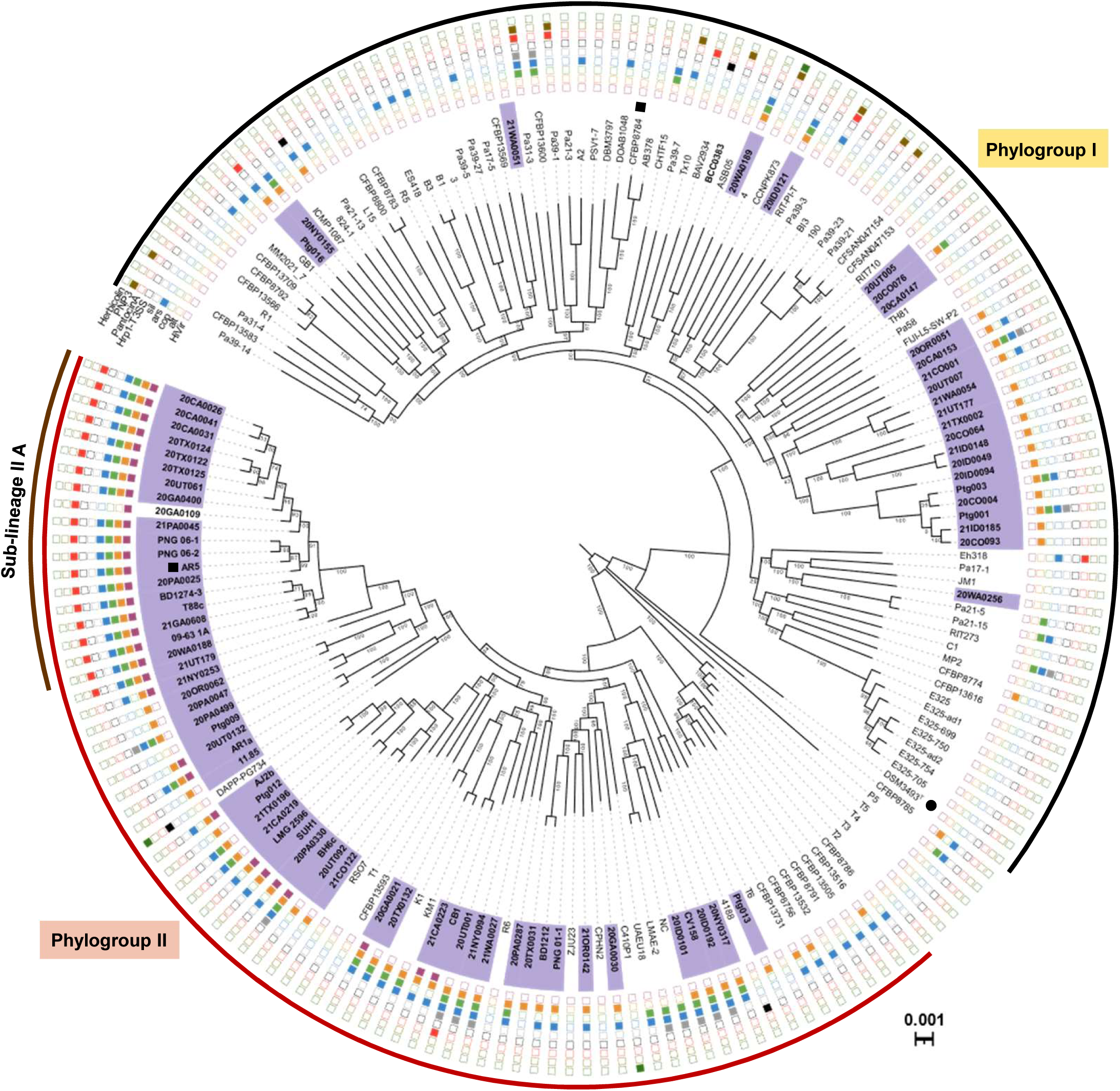
*P. agglomerans* strains divided into two well represented Phylogroups. Approximately-maximum-likelihood phylogenetic tree constructed using FastTree v.2.1.11 based on the 3,111 core genes shared among 181 *Pantoea agglomerans* strains. The tree was constructed using the general time reversible (GTR) nucleotide substitution model, and reliability of the branch groupings was estimated using the approximate likelihood-ratio test (aLRT). The resulting tree was visualized and edited in iTOL (v. 6.8, available at https://itol.embl.de/). The strain names are highlighted in bold to denote genomes sequenced in this study, and their association with onion is indicated in purple. The distributions of the virulence (HiVir and Hrp1-T3SS), antimicrobial (Pantocin A, PNP3, Herbicolin) and antimicrobial resistance (allicin (*alt*), copper (*cop*), arsenic (*ars*), silver (*sil*)) genes are indicated by the colored boxes next to bacterial strain names. The type strain of *P. agglomerans* DSM3493^T^ = LMG1286^T^ is indicated by the solid dot (●) next to the strain name. Proposed phylotype strains AR5 and CFBP8784 are marked with black squares. (▪). The two phylogroups are indicated by black (Phylogroup I) and red (Phylogroup II) outermost curved lines. CFBP8785, T2, T3, T4, T5 and P5 were not assigned to phylogroups.

Other analyses also supported the presence of two well-represented *P. agglomerans* phylogroups. The correlation plot of pairwise ANI values supported the same phylogroup assignments with 98% similarity shared within a phylogroup and 97% similarity between phylogroups. ANI values also excluded the six outgrouping *P. agglomerans* strains (Supplemental Figure S2 and Supplemental Table S10). The same strain groupings were also evident from the Roary gene presence/absence dendrogram. The maximum-likelihood phylogeny of a partial *gyrB* gene region previously validated for species discrimination was also largely congruent with *P. agglomerans* phylogroup assignments (Supplemental Figure S3). Thus, *gyrB* PCR assay and sequencing provided a simple method for *P. agglomerans* phylotyping for non-genome sequenced strains. The six out-grouping *P. agglomerans* strains showed inconsistent phylogroup placements in both the gene presence/absence dendrograms and the partial *gyrB*-sequence-based phylogeny relative to the core genome phylogeny or ANI analysis, which grouped them separately.

### Phylogenetic distribution of *P. agglomerans* onion strains and characterized *P. agglomerans* functional traits

Sequenced *P. agglomerans* strains isolated from onions fall into both Phylogroup I and Phylogroup II, with 25 and 54 isolates, respectively (Fig. 1). In addition, we identified a sub-lineage of twenty closely-related onion isolates (sub-lineage II A) in Phylogroup II (Fig. 1). Sub-lineage II A isolates were recovered from across the United States and from South Africa. Sub-lineage II A includes PNG 06-1 and PNG 06-2, the first *P. agglomerans* strains reported as onion pathogens in the United States isolated from diseased onions in Georgia in 2006 [Edens et al. 2006]. We identified the HiVir gene cluster in 31 *P. agglomerans* genomes, exclusively from Phylogroup II strains, including all 20 strains in sub-lineage II A. Strain 20GA0109 from Carolina geranium is the sole non-onion isolate encoding HiVir identified in this study.

The *alt* cluster conferring thiosulfinate tolerance, was widely distributed in both Phylogroups I and II although this was not universal trait. Out of 79 *P. agglomerans* onion strain assemblies, 63 encoded an *alt* cluster, including the 30 onion isolates encoding HiVir. In addition, *alt* clusters were identified from eight non-onion strains, assemblies including two foodborne strains isolated from kimchi, five strains from a *Brassica* sp. seed or *Raphanus* sp. seed or flower, as well as the *Gypsophila* gall-forming strain *P. agglomerans* pv. *gypsophilae* 824-1 (Supplemental Table S9).

Among secondary-metabolite gene clusters with characterized roles in biocontrol interactions or the *hrp1* T3SS associated with gall-formation in *P. agglomerans* strains, both the herbicolin biosynthetic cluster and *hrp1* T3SS clusters were identified sporadically among the 181 genomes, but not in any *P. agglomerans* onion genomes. The PNP3 biosynthetic cluster was identified in 10 Phylogroup I strains but was likewise, not in onion strain genomes. Conversely, the cluster for synthesis of pantocin A, a RiPP secondary metabolite, was identified in six Phylogroup I strains, including the onion strain Ptg016, and 21 onion strains Phylogroup II, including all 20 strains in sub-lineage II A (Fig. 1).

### The role of HiVir in *P. agglomerans* onion necrosis symptoms

We tested a subset of 18 onion-isolates (8 from Phylogroup I, 10 from Phylogroup II) isolated from nine different states in the U.S.A. for onion foliar, bulb, and RSN. We additionally included the *P. agglomerans* DSM 3494^T^ type strain (HiVir-negative) and an in-frame *hvrA* deletion mutant strain derivative of *P. agglomerans* AR1a. As has been observed in other *Pantoea* spp., the strains that possess the HiVir gene cluster could cause rapid necrosis in both onion leaf and bulb tissues [Asselin et al. 2018; Stice et al. 2020; Shin et al. 2023]. Conversely, no necrosis was observed in leaves or bulbs inoculated with HiVir-negative or in-frame *hvrA* deletion mutant strains (see Fig. 2A).

**Figure 2.**
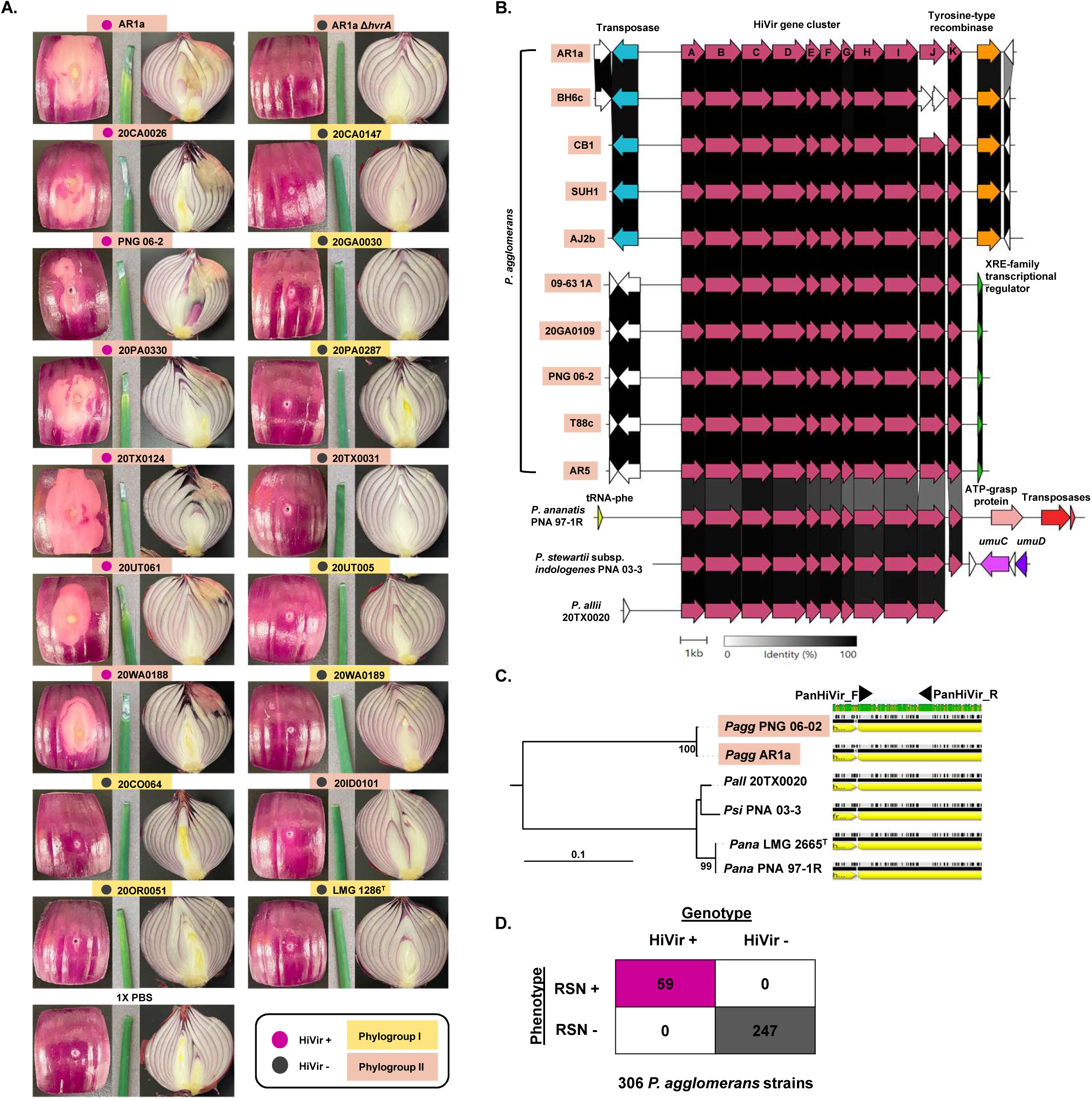
HiVir is the key onion necrosis inducing factor in *Pantoea agglomerans*. (**A**) Pathogenicity of HiVir-positive and -negative *P. agglomerans* strains was tested on onion, including leaves, detached bulb scales and whole bulbs. Onion necrosis was only observed in tissues inoculated with HiVir-positive strains, and symptoms were recorded 4 days post-inoculation (dpi) for scales and leaves, while bulbs were examined 1 week post-inoculation after incubation of the bulbs at 28°C. (**B**) A clinker diagram showing that the HiVir gene cluster structure is largely conserved across *Pantoea* species, with the degree of shading of lines between gene arrows representing nucleotide identity. *P. agglomerans*-HiVir is plasmid-borne and is found in two different plasmid contexts. (**C**) A maximum-likelihood phylogenetic analysis of the *hvrD* gene from various *Pantoea* species shows that *P. agglomerans*-*hvrD* forms a distinct group compared to that of *P. ananatis*, *P. allii*, and *P. stewartii* subsp. *indologenes*. Primers for detecting HiVir (PanHiVir) were designed based on conserved regions of the *hvrD* gene shared by at least four *Pantoea* species. (**D**) Screening 306 *P. agglomerans* strains using genotypic (PanHiVir-PCR assay) and phenotypic (red onion scale necrosis assay) methods revealed a 100% correlation between the presence of HiVir and pathogenicity on onion tissue. The strain names are highlighted according to phylogroups (yellow: Phylogroup I, pink: Phylogroup II).

Comparative analysis of the HiVir gene cluster across different *Pantoea* species revealed distinct genomic contexts in which the HiVir is encoded in different *Pantoea* species. By examining the genes flanking the HiVir, we identified two different plasmid contexts for the *P. agglomerans* HiVir in the closed genomes (Fig. 2B). Although the HiVir exhibited nearly identical nucleotide sequences between *P. agglomerans* strains, HiVir was notably different in sequence between *P. agglomerans* and *P. ananatis* strains. We observed 76% nucleotide identity between the HiVir gene clusters of *P. agglomerans* and that of *P. ananatis* 97-1R. The HiVir gene clusters of *P. allii* 20TX0020 and *P. stewartii* subsp. *indologenes* pv. *cepacicola* PNA 03-3 were more similar to that of *P. ananatis*, sharing 94 and 95% nucleotide identity, respectively (Fig. 2C, Supplemental Table S11).

The comparison of nucleotide sequences of HiVir gene clusters from four *Pantoea* species revealed that the genes *hvrA* to *hvrF* were the most conserved among species (Fig. 2B). This observation aligns with the previous discovery that the *hvrA* to *hvrF* genes are essential for inducing onion necrosis mediated by HiVir in *P. ananatis* and sufficient for synthesis of pantaphos [Shin et al. 2023; Polidore et al. 2024]. Conserved regions of *hvrD* were targeted for design of *Pantoea*-universal HiVir detection primers, referred to as PanHiVir (Fig. 2C).

For the 306 *P. agglomerans* strains, the majority collected as part of our multi-state survey, there was a perfect correlation between the presence of the *P. agglomerans* HiVir genotype (based on either existing genomic sequences or running the PanHiVir PCR assay) and a positive RSN phenotype (Fig. 2D). Of the 247 *P. agglomerans* strains lacking HiVir, none was capable of causing RSN symptoms. In contrast, all 59 strains that tested positive for HiVir exhibited RSN symptoms. All 59 HiVir positive strains were identified as Phylogroup II strains using these previously described methods for phylotyping. These findings provide additional evidence supporting the role of HiVir as a crucial and predictive factor responsible for pathogenicity of *P. agglomerans* strains to onion and its primary occurrence in Phylogroup II strains.

### Identification of *P. agglomerans* plasmid clusters, their distributions, and associated traits

The pool of 35 high-quality closed genomes (Supplemental Table S14) consisted of 35 chromosomes plus 125 plasmids for a total of 160 replicons. The chromosomes ranged from 3,978,822 to 4,410,564 bp; the plasmids ranged from 2,550 to 613,013 bp. A list of replicons and their statistics can be found in Supplemental Table S15. A summary of the replicon-level pangenome of each plaster group, determined using Roary, can be found in Table 1, and a complete list of the members of the “pangenomes” for each plaster group are shown in Supplemental Tables S18, S19, S20, S21, and S22. The analysis identified 31 plasmid clusters, or “plasters”, 9 of which contained greater than one member. A list of all plasters can be found in Supplemental Table S16. Supplemental Figure S6 shows as a heatmap the similarity of all pairs of chromosomes and plasmids. Hierarchical clustering was used to reorder the rows and columns to group similar pairs. The largest plaster, in terms of the combined length of members, contains the chromosomes of all 35 strains. Supplemental Figure S7 shows, as a heatmap, the similarity of all pairs of plasmids of this plaster. Supplemental Figure S8 shows, as a heatmap, the similarity of all pairs of LPP-1 plasmids. The second largest plaster contains a single plasmid from each genome. These plasmids range from 511,735 to 613,013 bp (mean 554,533 bp) and contain genes for carotenoid biosynthesis [Choi et al. 2020]. Consistent with the observations and nomenclature of De Maayer et al. [2012], we denoted this plaster and these plasmids as “LPP-1”, or Large Pantoea Plasmids.

Two small plasters included plasmids from four strains (CFSAN047153, CFSAN047154, FC61912-B, and ROTS050421) that were all, coincidently, isolated in New York State from non-onion sources, and the Hrp1 T3SS-encoding pPATHpab and pPATH plasmids from strains *P. agglomerans* pv. *betae* 4188 and *P. agglomerans* pv. *gypsophilae* 824-1, respectively [Nissan et al. 2019; Geraffi et al. 2023]. We denote these plasters as “pENY” (<u>p</u>lasmid from <u>E</u>nvironmental <u>N</u>ew <u>Y</u>ork strains) (minimum 203,713 bp, maximum 207,534 bp, mean 205,443 bp) and “pPATH” (minimum 131,449 bp, maximum 156,057 bp, mean 143,753 bp), respectively. In the absence of other biological context and limited representatives, we left the remaining plasters unnamed.

The third largest plaster also contains a single plasmid from each *P. agglomerans* genome. These plasmids range from 143,524 to 211,326 bp. A separate plaster analysis of all closed *Pantoea* genomes from RefSeq showed that this plaster is specific to *P. agglomerans* genomes (data not shown). We denote this plaster and these plasmids as “pAggl”. Thus, *P. agglomerans* can be expected to have a minimum of three genomic replicons, the chromosome, as well as LPP-1 and pAggl plasmids. Roary analysis of pAggl plasmids identified a set of 70+ pAggl core genes. The pAggl core genome included genes with annotations suggesting roles in plasmid replication and partition, a toxin-antitoxin system, sensing of environment and signaling, transcriptional regulation, iron transport, lipopolysaccharide (LPS) modification, acetoin biosynthesis, and sugar transport and metabolism. These included the seven-gene *arn* cluster for synthesis and LPS modification with 4-amino-4-deoxy-l-arabinose. In addition, the HiVir gene cluster was encoded on pAggl plasmids in strains 09-63_1A, 20GA0109, AR5, PNG_06-02, and T88c, all of which are members of Phylogroup II sub-lineage II A.

The plasmids of fourth largest plaster were found in *P. agglomerans* onion isolates with the sole exception of pPAG04 from *P. agglomerans* pv. *gypsophilae* 824-1. We denoted this plaster and these plasmids as “pOnion”, reflecting its common association with *P. agglomerans* onion strains. Within the *P. agglomerans* onion strains used for plaster analysis, the *alt* gene clusters were only found on pOnion plaster plasmids. The HiVir gene cluster was encoded on the pOnion plasmid in five of closed genome strains of *P. agglomerans* in Phylogroup II. In addition, pOnion plasmids often encoded genes for tolerance to copper (7/15 pOnion plasmids) and other metals described below. Some sequenced strains carried two plasmids that fell within this plaster. Unlike plasmids in the pLPP1 and pAggl plasters, plasmids in the pOnion plaster were mosaic genetically, making it challenging to interpret their relationships. This mosaic nature was reflected in the network representations of the MASH-based plasmid clustering (Fig. 5C). The LPP-1 and pAggl plasmids formed tight and self-contained networks compared with the diffuse and open network of the pOnion plasmids. In addition, by adjusting the plaster analysis thresholds, we resolved three groupings of pOnion plasmids, A-, B-, and C-type. Using Roary to identify genes shared between plasmid groups, A-type and B-type pOnion plasmids did not share core genes. However, both shared different genes sets in common with the C-type plasmids. Using BRIG [Alikhan et al. 2011], some single pOnion plasmids were observed to carry large contiguous genetic regions corresponding with those present on each of the two plasmids found in dual pOnion strains. The core B-type and C-type plasmid genes included the *alt* gene cluster and other genes also present in Onion Virulence Region A of the *P. ananatis* pOVR plasmid [Stice et al. 2018] (Figure 5D). When the pPAG04 plasmid from *P. agglomerans* pv. *gypsophilae* 824-1 was removed from the analysis, the B-type and C-type shared genes that included plasmid replication and partitioning genes (Figure 5D). Initially, no core A-type genes were identified. However, when the 20CO076 plasmid D was removed from analysis, a set of core A-type and C-type genes was identified that includes *dsbD* disulfide reductase domain genes, *dsbG* disulfide isomerase and genes for other thiol/redox associated proteins (Figure 5B,D). Using BRIG and BLAST analysis, we observed that some single C-type pOnion plasmids carried large contiguous genetic regions corresponding with those present on each of the two plasmids found in dual pOnion strains A-type and B-type pOnion plasmids (Fig. GINA5D).

### Copper tolerance gene clusters in *P. agglomerans* plasmids

The *copABCDRS* copper tolerance gene clusters were found in the genomes of 53 of 79 *P. agglomerans* onion strains. The *cop* genes were encoded on pOnion plasmids in seven closed genomes or on the same contigs as *alt* in 32 of the draft genomes (data not shown). The *P. agglomerans* copper tolerance clusters in the onion strains could be divided into three types based on their co-association with gene clusters for arsenic tolerance (also divided into two types based on sequence similarities) and for silver tolerance (Fig. 1 and Fig. 3A). The *copABCDRS* copper tolerance genes were present in all three cluster types but displayed a minimum of 20% nucleotide sequence variation among groups. In the Type I cluster, (the most common in the genome assemblies), the copper tolerance gene clusters were co-associated with arsenic tolerance genes. In the Type II cluster, the arsenic, copper, and silver tolerance gene clusters were co-associated. In the Type III cluster, only the copper tolerance genes were present.

**Figure 3.**
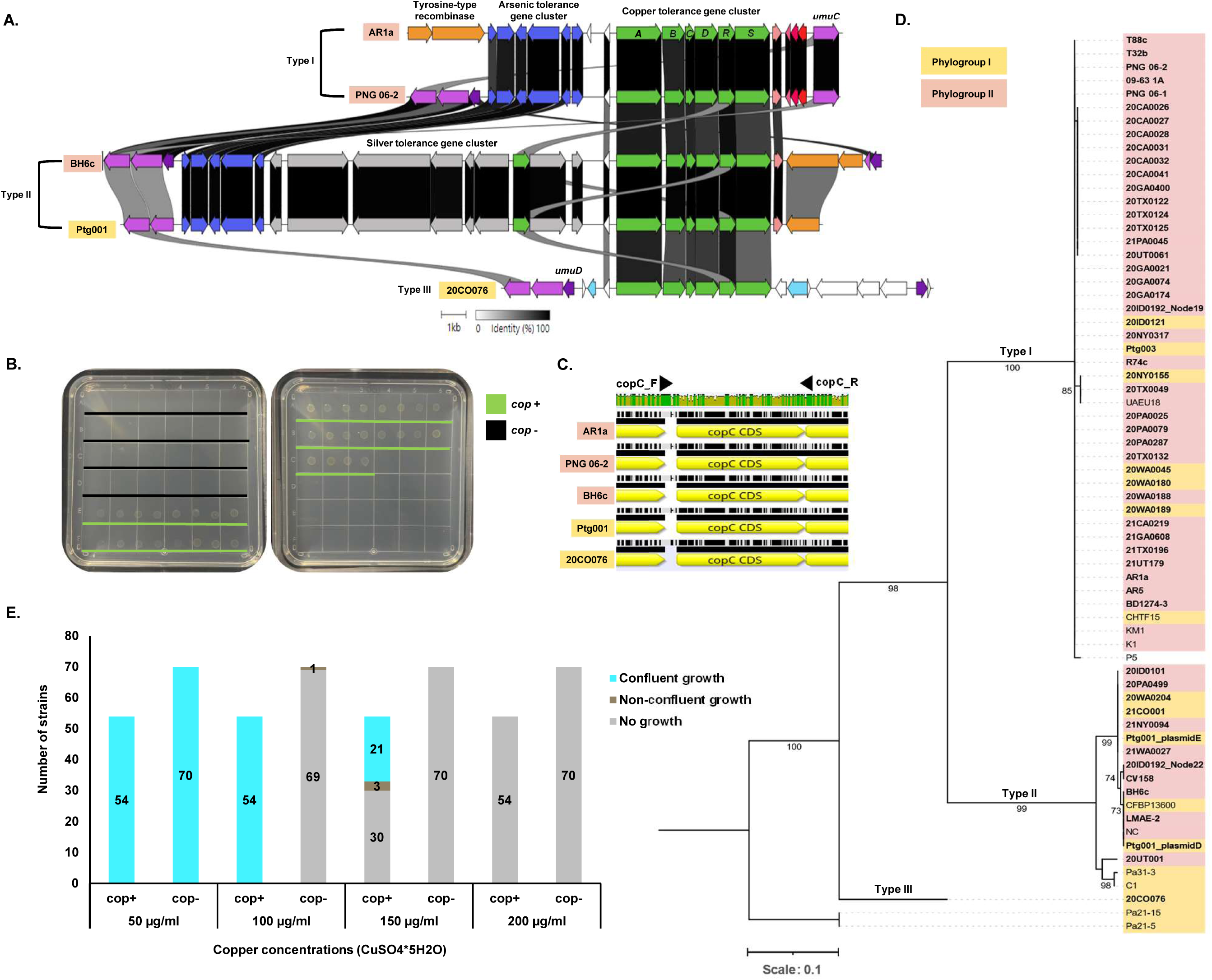
Identification of copper tolerance gene clusters in *Panotea agglomerans*. (**A**) Three distinct copper tolerance gene clusters were identified in *P. agglomerans* based on their nucleotide similarity and co-association of neighboring gene clusters annotated for arsenic and silver tolerance. (**B**) *P. agglomerans* strains possessing any type of copper tolerance gene cluster formed confluent colonies on CYE agar medium amended with 100 µg/ml copper sulfate, while strains lacking the copper tolerance gene cluster exhibited no growth or non-confluent growth. (**C**) Primers for detecting copper tolerance gene clusters were designed based on conserved regions flanking the *copC* gene. (**D**) A maximum-likelihood phylogenetic tree constructed using *copC* gene sequences extracted from available genome assemblies as well those obtained from PCR assays and sequencing using *copC* primers revealed three major groupings consistent with the types of copper tolerance gene clusters identified. (**E**) Screening 123 *P. agglomerans* strains using genotypic (copC-PCR assay) and phenotypic (copper sensitivity assay) methods confirmed that copper-tolerant strains (those with the copper tolerance gene cluster) can be distinguished from copper-sensitive strains at a concentration of 100 µg/ml copper sulfate.

Based on conserved regions flanking the *copC* gene, we designed PCR primers for identification of *copABCDRS* encoding strains (cop-positive) and sequence analysis of *copC* (Fig. 3C). Phylogenetic analysis using a maximum-likelihood approach revealed that for 65 *P. agglomerans copC* gene sequences extracted from genome assemblies or detected by the PCR assay and sequencing, the *copC* gene could be categorized into at least three distinct clades congruent with the copper tolerance gene typing scheme (Fig. 3D). Therefore, the *copC* gene can serve as a valuable marker not only for detecting *cop* gene presence, but also for determining the specific type of copper tolerance gene cluster present in a strain.

Based on phenotypic measurements of the relative copper sensitivity of 123 *P. agglomerans* strains (81 WGS strains and 42 strains genotyped with the *copC* PCR assay) (Supplemental Table S25), there was a consistent phenotypic difference in copper sensitivity thresholds, with all 54 *cop*-positive strains producing confluent growth in CYE medium augmented with 100 µg/ml copper sulfate while none of the 70 *cop*-negative strains produced confluent growth under the same conditions (Fig. 3B and 3E). Twenty one of the54 *cop*-positive strains, 21 displayed tolerance up to 150 µg/ml of copper sulfate (Fig. 3E). All three cluster types were represented among these relatively highly tolerant strains. This suggests that the specific copper tolerance gene cluster types are not predictive of relative copper tolerance.

### *P. agglomerans* CB1 pCB1C is a mobilizable onion pathogenicity plasmid

Using MacSyFinder (Supplemental Table S17) to predict the presence of features associated with plasmid transfer systems within our set of analyzed *P. agglomerans* replicons. 26 replicons were found to contain one or more likely mobilizable elements and 29 replicons contained conjugative T4SS’s of two types: F and G. Type-F was found on pOnion, pENY, and several other unclassified plasmids. Type-G was found on chromosomes of four *P. agglomerans* strains and on unclassified plasmids in six strains.

Notably, no LPP-1 nor pAggl plasmids encoded plasmid transfer systems and 10 pOnion plasmids carried a predicted intact conjugative Type-F T4SS [Cury et al. 2017]. Within this subset, five pOnion plasmids also encoded *alt* genes. Of these, pCB1C was also the only replicon from a strain with a closed genome to have both a HiVir cluster and putative intact T4SS cluster. Plasmid pCB1C appeared to have everything required to act as a transmissible pathogenicity plasmid: the HiVir cluster encoding the pantaphos toxin, the *alt* cluster protecting bacterial fitness after toxin deployment in onion, and an intact T4SS cluster encoding the machinery to move between strains.

Allelic exchange was used to create kanamycin resistance-marked derivatives of pCB1C of *P. agglomerans* CB1. *P. agglomerans* CB1 with the Kan^R^-marked pCB1C-AKan (Figure 4A) or pCB1C-BKan as donor strains were co-cultured with spontaneous rifampicin-resistant derivatives of four different *P. agglomerans* recipient strains lacking HiVir. Candidate exconjugant colonies from each mating, where available, were tested by PCR to confirm that the recovered colonies were recipient strains that had acquired pCB1C-AKan or pCB1C-BKan. PCR identified at least one putative exconjugant colony for each recipient strain, except for Pag J22c Rp^r^. A representative exconjugant from each of the matings of MMD61212-C Rp^r^, AR8b Rp^r^, and FC61912-B Rp^r^ with CB1 pCB1C-AKan were sequenced by ONT confirming that each strain retained their native plasmids while gaining pCB1C-Akan, except the AR8b Rp^r^ pCB1C-AKan exconjugant, which lost its native B-type pONION plasmid. (Figure 4d)

**Figure 4.**
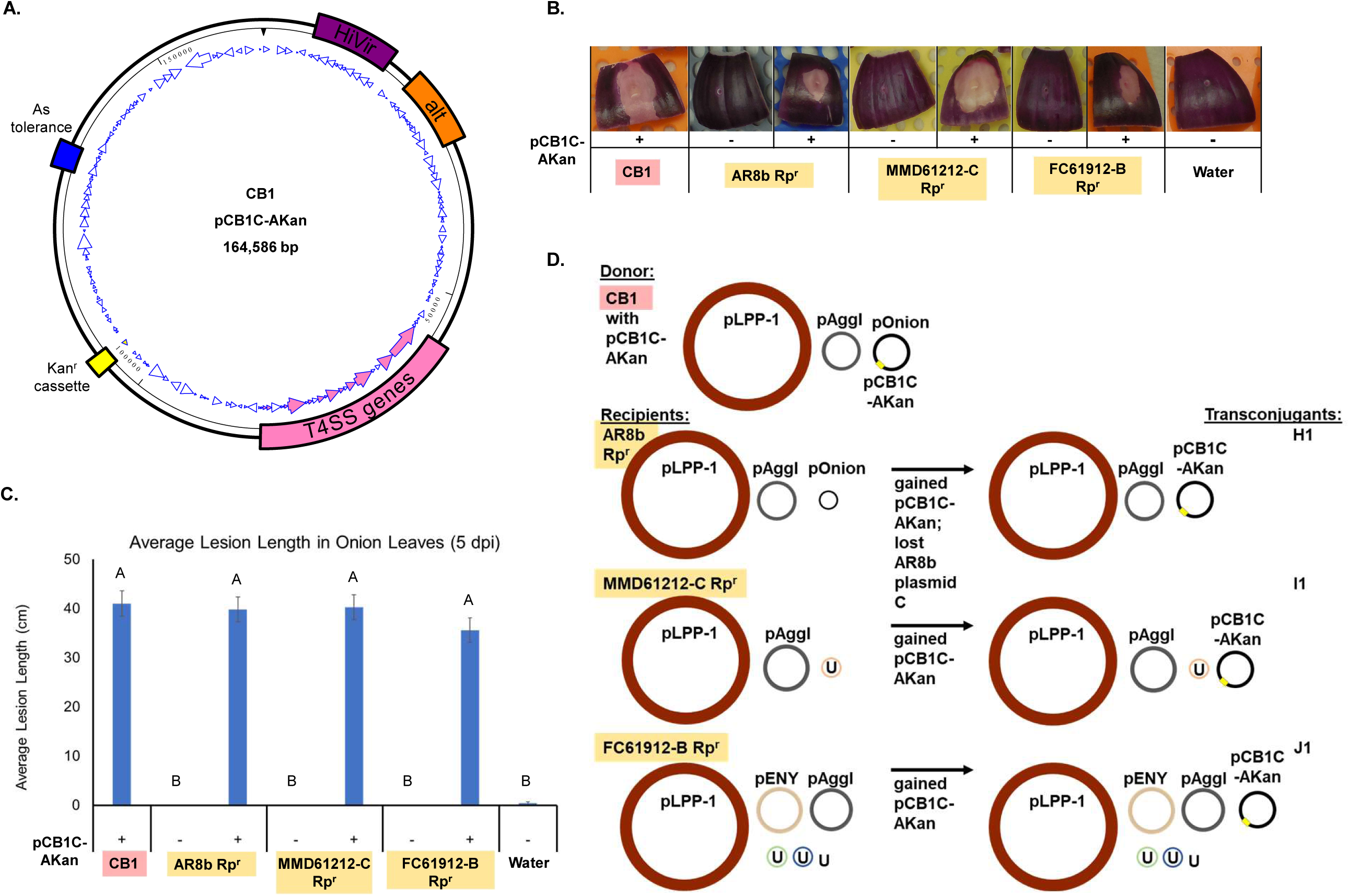
*Pantoea agglomerans* CB1 contains a mobilizable onion pathogenicity plasmid. (**A**) Representation of pCB1C-AKan, a version of pCB1C of strain CB1 modified to contain a kanamycin resistance gene cassette to facilitate tracking plasmid movement. pCB1C-AKan and pCB1C harbor the HiVir gene cluster, *alt* gene cluster, arsenic resistance genes, and a type IV secretion system gene cluster. The plasmid genome was visualized in SeqBuilder Version 17.3 (DNASTAR, Madison, WI). (**B**) *P. agglomerans* strains that received pCB1C-AKan (transconjugants) caused positive red scale necrosis resuls on detached fleshy scales from red onion bulbs. (**C**) Transconjugants were rendered pathogenic in onion leaves. (**D**) Whole genome sequencing revealed that recipient strain H1, derived from the non-pathogenic onion isolate *P. agglomerans* AR8b Rpr, lost its existing pOnion plasmid while adding pCB1C-AKan. Strains I1 and J1, derived from environmental isolates MMD61212-C Rpr and FC61912-B Rpr (not pathogenic to onions), added pCB1C-AKan without plasmid loss. Only plasmids (not chromosomal DNA) are depicted for the strains. Plaster category or name is listed for each plasmid. “U” (for unique) indicates that a plasmid did not form a plaster group with other plasmids in the study. pCB1C-AKan is a pOnion B type plasmid.

Overnight co-incubation of donor CB1 harboring pCB1C-AKan with recipient MMD61212-C Rp^r^ on water agar resulted in 2.3-4.7 x 10^-7^ transconjugants per initial donor cell or 2.8-4.1 x 10^-7^ transconjugants per initial recipient cell. There was also a 3.5-3.9-fold increase of the recipient strain and 3.6-5.6-fold increase of the donor strain on water agar over the course of incubation.

Three transconjugant strains tested, H1 (AR8b Rp^r^ pCB1C-AKan), I1 (MMD61212-C Rp^r^ pCB1C-AKan), and J1 (FC61912-B Rp^r^ pCB1C-AKan), caused a positive RSN response in each of three replicate onion scale pieces, consistent with bacteria producing pantaphos from the HiVir clusters.

In general, leaves inoculated with CB1 harboring the kanamycin-resistance cassette containing plasmid, and leaves inoculated with each of the transconjugants, developed lesions over five days. None of the original strains used as recipients caused lesions on the leaves. Two of 32 leaves treated with water (control treatment) developed lesions over the course of the experiment, indicating the presence of bacteria naturally occurring on the leaves. Details of the statistical analyses can be found in Supplemental Document S1.

## Discussion

### HiVir is a key onion pathogenicity factor in *P. agglomerans*

In our study of the genomic and phenotypic diversity of *P. agglomerans* strains associated with onion, we identified the HiVir gene cluster as a key factor for onion pathogenicity. The primary role of the HiVir in enabling strains of *P. ananatis* to cause symptoms on onion has been established recently [Asselin et al. 2018] and, as demonstrated in *P. ananatis*, HiVir-positive *P. agglomerans* strains are able to cause rapid necrosis in whole onion bulbs, detached fleshy scales and foliage. We observed perfect correlation between onion red scale necrosis phenotypes and HiVir gene cluster presence/absence across 306 *P. agglomerans* strains originating from the United States and internationally. Although both Phylogroup I and II strains of *P. agglomerans* were isolated from onion, all 59 HiVir-positive *P. agglomerans* onion isolates were identified as members of Phylogroup II.

Unlike the chromosomally encoded *P. ananatis* HiVir, the *P. agglomerans* HiVir is plasmid-borne and, in at least one case demonstrated to be plasmid-transmissible. We identified the *P. agglomerans* HiVir gene cluster in two distinct classes of plasmid, the conserved pAggl plasmids and the mosaic pOnion plasmids. The 20 pAggl-encoded HiVir strains comprise a closely related sub-lineage of Phylogroup II strains (sublineage II A) isolated from onions in South Africa and across the United States, with the first representative strain isolated in 2006 in Georgia State [Edens et al. 2006]. the oldest HiVir-positive strains from this study are pOnion-encoded HiVir strains isolated in 1977 in South Africa [Hattingh and Walters 1981]. In addition, although we identified no HiVir-positive Phylogroup I strains of *P. agglomerans*, we demonstrated that pCB1C is a mobilizable pOnion HiVir-encoding pathogenicity plasmid able to confer pathogenicity to onion to multiple non-pathogenic Phylogroup I strains of *P. agglomerans,* including environmental isolates.

The *P. agglomerans* HiVir genes were nearly identical in sequence among strains, regardless of their plasmid contexts. However, the *P. agglomerans* HiVir sequences were notably divergent from the HiVir sequences of three other *Pantoea* species, implying independent acquisition or divergence of *P. agglomerans* HiVir gene cluster. This is in contrast to the *alt* genes for thiosulfinate tolerance, which were nearly identical across the four *Pantoea* species. Consistent with their essential role in facilitating necrosis of onion tissues, the *hvrA-hvrF* genes displayed higher sequence conservation among *Pantoea* species than the necrosis non-essential genes *hvrG-hvrJ* between *P. agglomerans* and *P. ananatis* HiVir gene clusters. *P. agglomerans* BH6c, which has a RSN positive phenotype, carries a premature stop codon in its *hvrJ* gene. Instead *hvrJ* encoding a 336 amino acid protein, in BH6c there are two predicted genes encoding 180 and 150 amino acids in the same reading frame. The *hvrJ* gene was found previously to be non-essential but strongly contributing to onion necrosis in *P. ananatis* [Shin et al. 2023]. As the specific function of HrvJ has not been determined, it is unclear whether the truncated open reading frames (ORFs) encode functional proteins [Polidore et al. 2024].

### Strains from distinct *P. agglomerans* phylogroups were isolated commonly from onions

In agreement with previous work, we observed that *P. agglomerans* strains to be divided into at least two well represented phylogroups that are supported by core genome phylogeny, ANI, accessory gene distribution, and *gyrB* single locus sequence analysis. We did not see strong geographic signatures in the locations of onion strains in Phylogroups I and II, which is consistent with the routine movement of onion seed, bare-root seedlings, and onion sets across the United States for planting. Based on available whole genome sequences, Phlyogroup I strains have been isolated more frequently than Phylogroup II strains. However, members of Phylogroups I and II have been isolated from similar and overlapping niches, predominantly soil and plant sources including onion. In addition, both Phylogroup I and II onion strains typically carry the onion-niche-associated pOnion plasmids, implying some degree of genetic exchange among strains from the two Phylogroups. Based on similar geography, niche-association, and apparent genetic exchange, it is not clear what factors have driven phylogroup divergence. The six *P. agglomerans* strains that fell outside of Phylogroups I and II were also isolated from soil and plant sources. Both core genome phylogeny and ANI analyses grouped these strains separately from those in Phylogroups I and II. However, *gyrB* phylotyping did not. Thus, in the absence of other genetic information, *gyrB* phylotyping can be used relatively confidently to exclude a specific *P. agglomerans* strain as being a member of Phylogroups I or II.

The *P. agglomerans* type strain DSM 3494^T^ is one of two human clinical isolates included in our analysis and branched basally to all other Phylogroup I strains. While this could reflect niche-driven selective divergence, detailed studies of *P. agglomerans* in clinical settings are insufficient to make such claims. In addition, strain Tx10, isolated from sputum of a cystic fibrosis patient, is also a Phylogroup I strain but clustered more closely with other Phylogroup I strains. Although both Phylogroups I and II strains have been isolated from diverse sources, plants and soils are the most common source of isolation. We propose CFBP8784 and AR5, isolated from radish seed and diseased onion, respectively, as *P. agglomerans* phylotype strains based on their typical isolation sources, average MASH distances from other phlyogroup strains, average number of orthologs shared with other phylogroup strains, and availability in public culture collections.

### *P. agglomerans* plasmids encode key niche-adaptative traits

Inferring the genetic relationships among plasmids can be challenging due, in part, to their high degree of genetic instability. In this study, we use MASH distances based on Kmer signatures to categorize plasmids into clusters called plasters. Using MASH distance is not novel to this work. A very similar method is used in the consensus genome sequence assembler, Trycycler [Wick 2020], to group sequences from different assemblies by unitig. Nevertheless, our use of plaster analysis to group and name plasmids across genomes does appear to be novel. Since this approach uses MASH distances, which are computed using rank-ordered lists of medium length *k*-mers (mashtree defaults to *k*=21), our plaster analysis is based primarily on statistics of the plasmid sequences and not the sequences themselves. This differs from other approaches to plasmid classification that use the presence of specific replication and partitioning genes [Carattoli et al. 2014; Huisman et al. 2021; Giménez et al. 2022] or more traditional sequence alignment [Redondo-Salvo et al. 2021], or other methods to determine whether individual contigs in draft genomes belong to plasmids or chromosomes (e.g., Andreopoulos et al. [2021]; Sielemann et al. [2022]).

The method used in this study is fast, because it uses MASH distances and produces expected results. The analysis grouped chromosome and LPP-1 replicon sequences into individual plaster groups. The analysis also assigned pPATHpab and pPATHpag from strains 4188 [Nissan et al. 2019] and 824-1 [Geraffi et al. 2023], to a single cluster. Using this approach, we identified a cluster of conserved plasmids exclusive to *P. agglomerans*, which we named “pAggl”.

The pAggl family of plasmids was first described by Sulja et al. [2022] and named Large Pantoea Plasmid 2 (LPP-2). Our plaster analysis shows that the *P. agglomerans* instances of LPP-2 (namely, DAPP-PG734 plasmid 3 [Sulja et al. 2022], and P10c plasmid pPag1 [Smits et al. 2015]) are contained within the pAggl plaster. However, the *P. vagans* instance of LPP-2 (namely, C9-1 plasmid pPag1 [Smits et al. 2011]) is not. Similar to species-specific LPP-1 families of plasmids found across the *Pantoea* genus, perhaps species-specific LPP-2 families can be found as well. This is beyond the scope of this paper, and we chose the name “pAggl” to reflect the limits of our findings.

The chromosomal replicons clustered into two groups that correspond exactly to the Phylogroups I and II discussed above (Supplemental Table S7). Since chromosomes are thought to be propagated vertically, the replicons of the LPP-1 plaster should show a similar structure, which they do (Supplemental Figure S8). Interestingly, the replicons of the pAggl plaster also show the same structure (Supplemental Figure S9), which suggested they are vertically propagated as well. To summarize, our observations support the hypotheses that strains of *P. agglomerans* contain a chromosome and at least two plasmids, LPP-1 and pAggl, which are all propagated vertically. Closing the genomes of more strains of *P. agglomerans* will support or refute this.

Replicons of this pOnion plaster were divided into, roughly three groups: A-type, B-type, and C-type, (Supplemental Table S11). Although the A-type and B-type pOnion plasmids shared no core genes, they both shared distinct sets of core genes with the C-type pOnion plasmids. In closed onion strain genomes, the *alt* gene cluster, which confers tolerance to *Allium* thiosulfinates and increased bacterial fitness in necrotized onion tissue, was a shared trait of B-type and C-type pOnion plasmids (5D). When the non-onion strain plasmid pPAG04 was excluded, the B-type and C-type plasmid core genes also included plasmid replication and partitioning genes, implying potential common ancestry. The shared cluster of genes between A-type and C-type pOnion plasmids is enriched in thiol/redox and disulfide-associated genes, including two genes with DsbD disulfide reductase domains, perisplasmic DsbG disulfide isomerase, Bhl bifunctional sulfur transferase/dioxygenase as well as other sulfur thiol/redox genes. In plant pathogenic species of *Agrobacterium* and *Xylella*, Bhl constributes to tolerance of sulfide stress (de Lira et al. 2018). Perhaps this gene cluster, which we have named SCALiOn (<u>S</u>hared <u>C A L</u>ocus in p<u>On</u>ion), also contributes to increased bacterial fitness under the thiol stress conditions of necrotized onion tissue. The A-type and B-type pOnion plasmids of strain AR1a (pAR1aC and pAR1aD) align with independent regions of the C-type pOnion plasmid of PNG06-2 (pPNG06-2C), recapitulating the majority of its sequence (Figure 5, Supplmental Figure S12). There are several possible explanations for these results. One, fusion of two ancestral versions of the dual A-type and B-type AR1a plasmids C and D fused to form an ancestral version of T88c plasmid C, the C-type pOnion plasmid of PNG06-2 (see, for example, Roman-Reyna et al. [2023]). An alternative hypothesis is that an ancestral version of T88c plasmid split to form ancestral versions of the AR1a plasmids C and D. Our results do not provide strong support for either hypothesis. We expect that the closing of more genomes will shed light into the historical relationship among the members of the pOnion plaster.

The pOnion plasmids display a high degree of genetic instability compared with LPP-1 and pAggl plasmids. Even within the closely related strains of sub-lineage II A, C-type pOnion plasmids varied in size, and strain 20GA0109, isolated from a Carolina geranium weed in an onion field, lacks a pOnion plasmid. A parsimonious explanation is that 20GA0109 lost its *alt-*encoding pOnion plasmid while colonizing the weed host.

**Figure 5.**
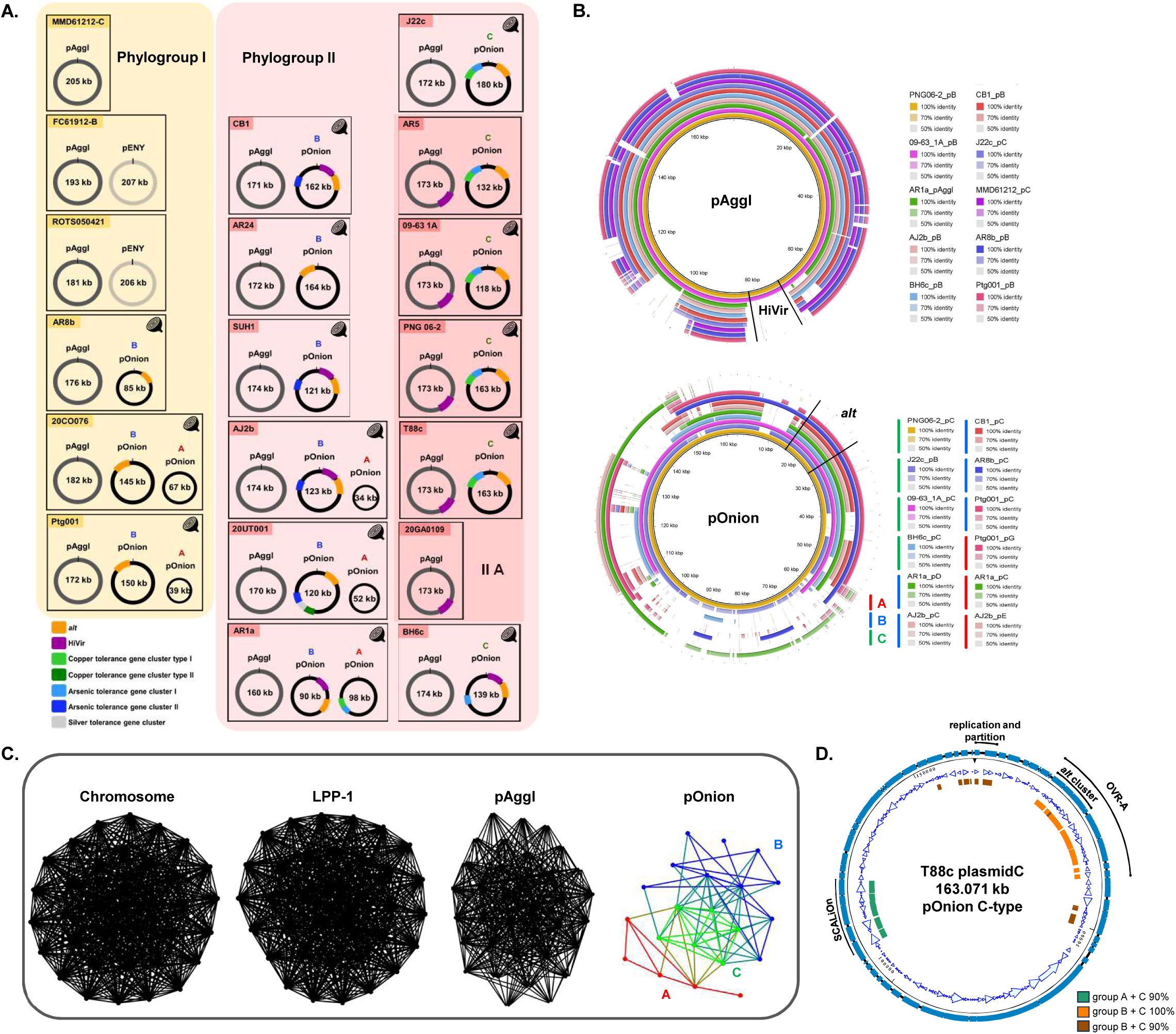
*Pantoea agglomerans* plasmids pAggl and pOnion encode onion virulence traits. (**A**) The selection of pAggl and pOnion from a subset of *P. agglomerans* strains from Phylogroup I (yellow) and II (pink) to illustrate the distribution of onion virulence (HiVir and *alt*) and metal (copper, arsenic, and silver) tolerance gene clusters found on both plasmids. The key for these traits is at the bottom of (A), and their positions on the plasmid relative to the *repA* gene are indicated. (**B**) BRIG [Alikhan et al. 2011] diagram showing a degree of genetic conservation between pAggl (conserved) and pOnion (mosaic) plasmid types (**C**) Network analysis based on K-mer statistics shows groupings of replicons found in 35 *P. agglomerans* strains with closed genomes. pOnion network nodes are colored: red = A-type, blue = B-type, green = C-Type. pOnion edges are colored: red = A-to-A, blue = B-to-B, green = C-to-C, other = cross group edges. The pAggl-network resembles those of the vertically transmitted replicons such as chromosome and Large Pantoea Plasmid (LPP-1) whereas pOnion shows an open network structure consistent with its mosaic nature. (**D**) A diagram of T88c plasmid C, a pOnion C-type plasmid, with colored boxes denoting genes (as determined by prokka annotation) conserved in >90% of type A+C plasmids (green), conserved in 100% of type B+C plasmids (orange), or conserved in >90% of type B+C plasmids (brown). No genes were conserved in 100% of type A+C plasmids. DNA was visualized in SeqBuilder Version 17.3 (DNASTAR, Madison, WI). Regions containing replication and partiton genes, *alt* and OVR-A gene clusters, and the SCALiOn region are labeled.

The messy structure of the pOnion plaster illustrates the impact of several design choices in the plaster analysis. First, the use of MASH distance, as opposed to sequence identity or homologous proteins, provides a simple, fast, purely computational measure of plasmids relationships. Second, the use of connected components, instead of approximate clique or other more traditional clustering methods, can produce “sloppy” clusters, which, as demonstrated by the pOnion cluster, can reveal complex relationships among plasmids that might not otherwise be apparent.

The use of plaster analysis in other studies may call for revisiting these choices. For instance, COPLA [Redondo-Salvo et al. 2021] (Classifier Of PLAsmids), provides a different graph-based approach to plasmid classification. COPLA uses ANI instead of MASH to quantify plasmid similarity. Also, COPLA uses Hierarchical Stochastic Block Modeling (HSBM) instead of connected components to collapse a similarity graph in to plasmid taxonomic units, or PTUs, which are roughly analogous to plasters defined in this study.

Shetty et al. [2023] proposed a plasmid classification that partitions plasmids of *Pantoea* into one of four categories: 1) plasmids that contribute to genetic diversification of *Pantoea*, 2) plasmids that contribute to the pathogenicity of *Pantoea*, 3) cryptic plasmids, and 4) plasmids that play a role in the metabolic versatility of the genus *Pantoea*. Plasmids are classified based upon their perceived role in fitness and environmental adaptation of strains in which they occur. This study also leverages the names of the plasmids assigned when the strains were deposited. This approach compares plasmids using an objective metric (MASH distance) and groups them using a clearly stated, albiet arbitrary, cutoff (MASH distance ≤0.05). Nevertheless, it is possible to interpret our results within their framework. For instance, plasmids assigned to our LPP-1 and pAggl contribute to genetic diversity of the bacterial strains, and pOnion and pPATH contribute to pathogenicity of strains to specific plant species. However, the divisions are not clear cut, as virulence factors, such as HiVir, can be found on pAggl as well as pOnion.

### Copper tolerance genes and their co-occurrence with *alt* thiosulfinate tolerance genes

We identified copper tolerance genes (*copABCDRS*) as a common antimicrobial resistance feature in onion-associated strains of *P. agglomerans* that resemble copper tolerance gene clusters previously characterized from both *Pseudomonas* and *Xanthomonas* plant pathogens [Rademacher and Masepohl 2012]. The *P. agglomerans cop* gene clusters could be divided into three types based on their associations with arsenic (*ars*) and silver (*sil*) tolerance genes and *copC* gene phylogeny. All three types of cop gene clusters were associated with similar phenotypic thresholds for copper tolerance *in vitro* (100 µg/ml).

A whole-genome-based search of *ars*, *cop,* and *sil* resistance gene clusters across 181 *P. agglomerans* strains revealed an unexpected co-association of *alt* and *cop* genes on the same contigs, particularly in onion-associated *P. agglomerans* strains (Fig. 1). For example, the genome-wide search identified 53 *cop*+ strains, of which 43 were alt+ and 10 were alt-. The *cop* genes were encoded on pOnion plasmids in seven strains with closed genomes, and on the same contig as the *alt* thiosulfinate tolerance genes in 27 draft genomes. While frequent co-occurrence of alt and *cop* gene clusters has been observed, co-occurrence of HiVir and *cop* on a single contig has only been noted in one strain 20GA0021, out of 181 *P. agglomerans* strain genomes. The co-occurrence of thiosulfinate tolerance and copper tolerance as well as other heavy metals in pONION plaster plasmids resembles multidrug-resistance plasmids. Prevalence of copper tolerance genes in *P. agglomerans* onion strains, often co-inherited alongside *alt* thiosulfinate tolerance genes, may accounted for the limited the efficacy of copper-based bactericides for management of onion center rot, but this has not been tested directly in the field. However, numerous field trials conducted to evaluate the efficacy of bactericides against bacterial bulb rot disease in onion have shown that copper-based bactericides are not effective in many trial locations [du Toit et al. 2022; Koirala et al. 2023].

### Plasmid-transmissibility of onion pathogenicity

Experiments with pCB1C-AKan, a kanamycin-resistant derivative of pCB1C, confirmed that the plasmid could transfer spontaneously between *P. agglomerans* strains under laboratory conditions and that acquisition of this plasmid could transform a non-pathogenic *P. agglomerans* strain into an onion pathogen. The onion strain AR8b pCB1C-AKan exconjugant, lost its endogenous pOnion plasmid, potentially due to plasmid incompatibility. In transconjugants of the environmental strains FC61912-B and MMD61212-C, which lacked native pOnion plaster plasmids, pCB1C-AKan was acquired without loss of native plasmids.

While pOnion plaster plasmids were found in both Phylogroup I and II, the HiVir-encoding variant pOnion plasmids were only identified among Phylogroup II strains. However, our results demonstrate that pCB1C-AKan can be transferred to a Phylogroup I strain and confer onion pathogenicity. It seems that there is no molecular barrier that explicitly prevents Phylogroup I strains from acquiring a pOnion-HiVir plasmid. In addition, Phylogroup I and II strains have both been isolated from onions, providing potential opportunities for plasmid exchange.

Nevertheless, other factors may affect the transmission, acquisition, and maintenance of pOnion plasmids by Phylogroup I and II strains in the environment. Onion strains from both phylogroups often carry pOnion plaster plasmids, which may be in the same incompatibility groups and limit exchange of pOnion plasmids. In addition, co-occurrence of a HiVir cluster and predicted intact T4SS cluster on the same plasmid appears to be rare, occurring only in pCB1C among the closed genomes evaluated in this study. However, examples of intact T4SS clusters in pOnion plasmids lacking HiVir were identified. Computationally-predicted, intact T4SS clusters were also commonly found on other *P. agglomerans* plasmids and on the chromosome, suggesting that many replicons have the potential for movement among *P. agglomerans* strains. Additionally, genes annotated as encoding integrases, transposases, and recombinases were common in pOnion plasmids and elsewhere in the genome. Given the mosaic nature of pOnion-group plasmids, genes involved in recombination likely have had a significant impact on the movement of virulence factor genes, including movement of genes from mobilizable plasmids to immobile replicons and vice versa. Within onion-pathogenic *P. agglomerans* strains, the HiVir gene cluster is highly conserved but can be found in the highly mosaic pOnion and the more conserved pAggl, demonstrating movement of this key pathogenicity factor between replicons.

## Conclusion

Draft and closed genomes of *P. agglomerans* described in this study will provide valuable resources for future research into bacterial and plasmid ecology, with potential agricultural impacts. Important questions remain unanswered. Why are *P. agglomerans* strains in Phylogroups I and II so commonly isolated from onion compared to *P. agglomerans* strains outside these groups? Are there differences in their survival in onions, their ability to be transmitted by insects such as thrips, or their ability to be seed borne? Copper tolerance genes, previously described in *P. agglomerans* [Tho et al. 2019; Corsini et al. 2016], were relatively common in strains isolated from onions and are often co-encoded with alt genes on plasmids. Does the presence of copper tolerant *P. agglomerans* strains in a field affect growers’ decisions to apply copper pesticides? There is evidence that the HiVir and *alt* gene clusters are mobilizable from some strains, but the distribution of HiVir in *P. agglomerans* strains appears limited, as this cluster was detected only in strains from one phylogroup in this study. Are there factors that limit the spread of HiVir? Are there other hosts or environments where the HiVir cluster has a significant benefit or fitness cost to *P. agglomerans*? While the answers to these and other questions may impact agricultural practices, research is needed to determine those potential impacts.

## Acknowledgments

We thank Dr. Erika Mudrak and Dr. Lynn Johnson of the Cornell Statistical Consulting Unit for help with data analysis.

We thank Benjamin Wood and Dr. James Woodhall, University of Idaho, Parma Research and Extension Center for providing isolates from Idaho and Eastern Oregon”.

This work was not possible without the help and support of the late Dr. Steven Beer. The contributions of Dr. Beer and his lab to bacterial onion disease research are enormous, and he generously provided many of the strains studied in the present work. Perhaps just as importantly, he connected the UGA and USDA labs and encouraged our collaboration. The paper would not have happened without him.

This work is supported by Specialty Crops Research Initiative Award 2019-51181-30013 from the USDA, National Institute of Food and Agriculture. Any opinions, findings, conclusions, or recommendations expressed in this publication are those of the author(s) and do not necessarily reflect the view of the U.S. Department of Agriculture.

The U.S. Department of Agriculture (USDA) is an equal opportunity provider and employer. Mention of trade names or commercial products in this publication is solely for the purpose of providing specific information and does not imply recommendation or endorsement by the USDA.

## Supplemental files

**Supplemental Figure S1.**

A heatmap visualization of the presence (pink) and absence (yellow) genes (n= 27,204) observed across 181 *P. agglomerans* strains. The dendrogram drawn based on the gene presence and absence categorizes *P. agglomerans* strains into two groups whose members are consistent with those of the Phylogroup I (indicated by black line) and Phylogroup II (red line).

**Supplemental Figure S2.**

The correlation plot, based on ANI (average nucleotide identity) values, categorizes *P. agglomerans* strains according to their phylogroups. Pairwise ANI values for 187 *P. agglomerans* strains were calculated using FastANI (v. 1.3.3) and organized through a hierarchical clustering method employing the Pearson correlation matrix. The strains are partitioned into two groups, denoted by red and blue, aligning with phylogroup distinctions (blue corresponds to Phylogroup I, while red corresponds to Phylogroup II).

**Supplemental Figure S3: The phylogeny of *P. agglomerans*, based on the *gyrB* gene is largely congruent with the core genome phylogeny.**

(**A**) An approximately-maximum-likelihood phylogenetic tree, constructed using FastTree v.2.1.11, relies on the 3111 core genes shared among 181 *P. agglomerans* strains (Figure 1). Maximum-likelihood phylogenetic trees, constructed based on partial *gyrB* gene sequences targeted by primers designed by (**B**) [Bonasera et al. 2014] and (**C**) [Brady et al. 2008], indicate that the Bonasera *gyrB* region reveals a phylogeny congruent with that of the core genome. Strains not assigned to phylogroups in the core genome tree (CFBP8785, T2, T3, T4, T5, and P5) form part of phylogroups in *gyrB* trees. Phylogroups are highlighted according to the phylogroup designation in the core genome tree, with a yellow box representing Phylogroup I and red representing Phylogroup II. The association of strains with onions is indicated by the purple coloring of strain names.

**Supplemental Figure S4.**

Extended phylogroup identification of 226 *Pantoea agglomerans* strains using maximum-likelihood phylogenetic analysis of partial *gyrB* gene as targeted by primers designed by [Bonasera et al. 2014]. *Tatumella ptyseos* ATCC33301^T^ was used as an outgroup, and branch lengths were hidden. Phylogroups are highlighted: Phylogroup I is marked with a yellow box, and Phylogroup II with pink.

**Supplemental Figure S5.**

A maximum-likelihood phylogenetic tree constructed based on nucleotide sequences of thiosulfinate tolerance gene (*alt*) clusters belonging to *P. agglomerans*, *P. ananatis*, and *P. allii* (Left). A bootstrap analysis of 1000 replicates was conducted to assess the reliability of groupings, and the tree was rooted at the midpoint. A clinker diagram shows the genetic arrangement of *alt* clusters (Right). Shaded lines between gene arrows represent nucleotide identity.

**Supplemental Figure S6.**

A combined heirarchical clustering and heatmap showing the relationship between the all of the replicons (chromosomes and plasmids) studied. The distances between replicons was computed using Mashtree v. 1.4.5 [Katz et al. 2019]. The figure was produced using R v. 4.3.1 [R Core Team 2023] with the ComplexHeatmap v. 2.14.0 library [Gu 2022]. Values shown are 100×(1-MASH), which is analogous to percentage identity.

**Supplemental Figure S7.**

A combined heirarchical clustering and heatmap showing the relationship between the **chromosomal** replicons studied. The distances between replicons was computed using Mashtree v. 1.4.5 [Katz et al. 2019]. The figure was produced using R v. 4.3.1 [R Core Team 2023] with the ComplexHeatmap v. 2.14.0 library [Gu 2022]. Values shown are 100×(1-MASH), which is analogous to percentage identity.

**Supplemental Figure S8.**

A combined heirarchical clustering and heatmap showing the relationship between the **LPP-1** replicons studied. The distances between replicons was computed using Mashtree v. 1.4.5 [Katz et al. 2019]. The figure was produced using R v. 4.3.1 [R Core Team 2023] with the ComplexHeatmap v. 2.14.0 library [Gu 2022]. Values shown are 100×(1-MASH), which is analogous to percentage identity.

**Supplemental Figure S9.**

A combined heirarchical clustering and heatmap showing the relationship between the **pAggl** replicons studied. The distances between replicons was computed using Mashtree v. 1.4.5 [Katz et al. 2019]. The figure was produced using R v. 4.3.1 [R Core Team 2023] with the ComplexHeatmap v. 2.14.0 library [Gu 2022]. Values shown are 100×(1-MASH), which is analogous to percentage identity.

**Supplemental Figure S10.**

A combined heirarchical clustering and heatmap showing the relationship between the **pOnion** replicons studied. The distances between replicons was computed using Mashtree v. 1.4.5 [Katz et al. 2019]. The figure was produced using R v. 4.3.1 [R Core Team 2023] with the ComplexHeatmap v. 2.14.0 library [Gu 2022]. Values shown are 100×(1-MASH), which is analogous to percentage identity.

**Supplemental Figure S11.**

A combined heirarchical clustering and heatmap showing the relationship between the **pENY** replicons studied. The distances between replicons was computed using Mashtree v. 1.4.5 [Katz et al. 2019]. The figure was produced using R v. 4.3.1 [R Core Team 2023] with the ComplexHeatmap v. 2.14.0 library [Gu 2022]. Values shown are 100×(1-MASH), which is analogous to percentage identity.

**Supplemental Figure S12.**

An alignment of AR1a’s plasmids C and D against T88c’s plasmid C. All three plasmids are members of the pOnion plaster. These results demonstrate the the sequence of T88c plasmid C can be reconstructed using sequence from AR1’a plasmids C and D. T88c plasmid C CDSs with possible roles in partition and replication are filled in black, genes with annotations suggesting involvement in recombination (e.g. transposases, integrases) are filled in purple, genes annotated as umuC or umuD are filled in green, and CDSs involved in type IV secretion are filled in pink. The figure was generated using the NCBI BLAST website, the blastn tool, the “Align two or more sequences” option, optimized for somewhat similar sequences. T88c plasmid C sequence was used as the query, and AR1a plasmids C and D as subject sequences. The graphical representation of the alignment was pasted below a linear representation of the plasmid generated in SeqBuilder 17.3 (DNAStar, Madison, WI) using pgap annotations.

**Supplemental Figure S13.**

This figure shows pAggl plaster plasmids pAR5_B and pAR1aB with core pAggl genes labeled with green boxes. Core genes for pAggl were determined using Roary based on prokka annotations. Arrows denote CDSs from RefSeq annotations. Some genes appeared to occur in clusters where genes had similar function or seemed to act in a single pathway. Where their annotations allowed reasonable guesses of function, gene clusters are labeled with yellow bars and expected functions.

**Supplemental Table S1.**

The 618 *Pantoea agglomerans* strains that are mentioned in this manuscript and the tables in which they are mentioned.

**Supplemental Table S2.**

A list of 502 *Pantoea* strains tested for red onion scale necrosis assay (RSN), and *P agglomerans*-specific and *Pantoea*-HiVir PCR assays.

**Supplemental Table S3.**

A comprehensive list of the primers utilized in this study for gene detection, sequencing, and genetic manipulations.

**Supplemental Table S4.**

QUAST quality assessment results for the 87 *Pantoea agglomerans* genomes sequenced in this study.

**Supplemental Table S5.**

A table displaying the BUSCO genome assembly and annotation completeness scores for 81 genomes sequenced in this study.

**Supplemental Table S6.**

A compilation of 125 *Pantoea agglomerans* strains subjected to whole genome sequencing (WGS) and *gyrB* gene sequencing in this study. Closed, complete genomes were generated for 19 strains, draft genomes were made for 68 strains, and *gyrB* gene sequences were made for 38 strains.

**Supplemental Table S7.**

A list of the 100 public *Pantoea* genomes employed in this study for genome analysis, detailing strain information including source, year, and place of isolation, and their respective GenBank accession numbers.

**Supplemental Table S8.**

Type strains of *Pantoea* species and their respective GenBank accession numbers from which the *gyrB* gene was extracted for phylogenetic analysis.

**Supplemental Table S9.**

Metadata information utilized for annotating the 181 *Pantoea agglomerans* strains that appear in Figure 1.

**Supplemental Table S10.**

A pairwise average nucleotide identity (ANI) comparison table of 187 *Pantoea agglomerans* genomes.

**Supplemental Table S11.**

A pairwise nucleotide identity comparison of HiVir gene clusters among *Pantoea* species.

**Supplemental Table S12.**

Results from preliminary conjugation experiments using *Pantoea agglomerans* strain CB1 harboring a version of the pCB1C plasmid with a kanamycin-resistance cassette as donor and four non-pathogenic strains of *P. agglomerans* as potential recipients. For each mating pair, three colonies, or fewer if three were not available, were screened via PCR with primers designed to amplify the donor strain (CB1 3125-F/3125-R), the recipient strains (ImpA F1/R2), or from within the HiVir cluster (Pag 3283 F1/R1), which is present on pCB1C but not in any of the wild-type recipient strains. The results are color coded. Green means the colonies gave the expected PCR result. Red means PCR indicated rifampicin-resistant donor colonies breaking through. Yellow means the PCR result was ambiguous. Putative transconjugants were selected for genome sequencing from conjugations with Pagg CB1 pCBC-AKan colony 30 as the donor. Putative transconjugants were recovered from conjugations with AR8b Rp^r^, MMD61212-C Rp^r^, and FC61912-B Rp^r^ as donors, but not with J22c Rp^r^.

**Supplemental Table S13.**

Details of strains used in conjugation experiments. The replicons found in each strain are listed with some relevant features. TXXScan and CONJScan results using MacSyFinder are listed for the strains. Strain derivatives (rifampicin resistant or altered to carry a kanamycin resistance cassette) are presumed to have identical results to wild-type strains. Phage predictions were generated with PHASTER [Arndt et al. 2016] and PHAST [Zhou et al. 2011].

**Supplemental Table S14.**

This table lists the strains whose genomes were used in the plaster analysis.

Initially, 38 closed, complete genomes were collected into a pool of *P. agglomerans* genomes. The table lists, for each strain, its “phylogroup” and GenBank “accession” identifier.

In order to confirm these taxonomic identifications, we computed the ANI scores of each strain compared to the *P. agglomerans* type strain (here denoted strain FDAARGOS1447). We found that the ANI scores of 3 strains (33.1, FL1, and HJS002), as found in the “fastani” column, fell below the typical species cutoff of 95% and were removed from the pool. We used BUSCO to evaluate the integrity of the remaining 35 assemblies; we found that that “busco_C” and “busco_D” scores ranged from 98.7%-99.5% and 0.2%-2%, respectively, indicating the quality of these assemblies ranged from good to excellent.

The “new” column records whether the genome assembly is new to this study. The “used” column records whether or not the genome was ultimately used as input for the plaster analysis.

**Supplemental Table S15.**

A list of replicons used in the plaster analysis and their statistics. The pool of 35 genomes listed in Supplemental Table S14 consisted of 35 chromosomes plus 125 plasmids for a total of 160 replicons. The chromosomes ranged in sizes from 3,978,822bp to 4,410,564; the plasmids ranged in sizes from 2,550bp to 613,013bp.

**Supplemental Table S16.**

A list of all the plaster assignments by our plaster analysis to the 160 replicons listed in Supplemental Table S15. Our analysis identified 31 distinct plasmid clusters, or “plasters”, 9 of which were non-trivial (>1 member). We assigned previously published names to 3 of the non-trivial plaster plasters (“chromosome”, “LPP-1”, and “pPATH”) and new names to 3 others (“pAggl”, “pOnion”, and “pENY”).

**Supplemental Table S17.**

**Predicted mobilizable and conjugative elements**

Mobilizable and conjugative elements were predicted by MacSyFinder v. 2.1.2 [Abby et al. 2014; Neron et al. 2022] using the CONJScan/Plasmids v. 2.0.1 models [Abby et al. 2016]. For each replicon, the number of predicted instances of each submodel is shown. The submodels with predicted include, MOB (mobilizable elements), T4SS_typeF and T4SS_typeG (functional conjugative T4SS’s), and dCONJ_typeG and dCONJ_typeT (decayed conjugative T4SS’s).

**Supplemental Table S18.**

**Pan-replicon Analysis Results of Chromosome Replicons**

A list of all of the gene groups identified by Roary [Page et al. 2015] for replicons belonging to the “chromosome” plaster. Each row reports a single gene group. The “Gene” and “Annotation” values report the Prokka [Seemann 2014] predicted gene name and gene product for the genes in the group. In the case that multiple different gene names or products are predicted, all predictions are listed. The “Count” and “Pers” columns report the number and percentage of replicons in the plaster group containing members of the genome group. The remaining columns indicate the presence of the gene group in individual replicons.

**Supplemental Table S19.**

**Pan-replicon Analysis Results of LPP-1 Replicons**

A list of all of the gene groups identified by Roary [Page et al. 2015] for replicons belonging to the “LPP-1” plaster. Each row reports a single gene group. The “Gene” and “Annotation” values report the Prokka [Seemann 2014] predicted gene name and gene product for the genes in the group. In the case that multiple different gene names or products are predicted, all predictions are listed. The “Count” and “Pers” columns report the number and percentage of replicons in the plaster group containing members of the genome group. The remaining columns indicate the presence of the gene group in individual replicons.

**Supplemental Table S20.**

**Pan-replicon Analysis Results of pAggl Replicons**

A list of all of the gene groups identified by Roary [Page et al. 2015] for replicons belonging to the “pAggl” plaster. Each row reports a single gene group. The “Gene” and “Annotation” values report the Prokka [Seemann 2014] predicted gene name and gene product for the genes in the group. In the case that multiple different gene names or products are predicted, all predictions are listed. The “Count” and “Pers” columns report the number and percentage of replicons in the plaster group containing members of the genome group. The remaining columns indicate the presence of the gene group in individual replicons.

**Supplemental Table S21.**

**Pan-replicon Analysis Results of pOnion Replicons**

A list of all of the gene groups identified by Roary [Page et al. 2015] for replicons belonging to the “pOnion” plaster. Each row reports a single gene group. The “Gene” and “Annotation” values report the Prokka [Seemann 2014] predicted gene name and gene product for the genes in the group. In the case that multiple different gene names or products are predicted, all predictions are listed. The “Count” and “Pers” columns report the number and percentage of replicons in the plaster group containing members of the genome group. The remaining columns indicate the presence of the gene group in individual replicons.

**Supplemental Table S22.**

**Pan-replicon Analysis Results of pENY Replicons**

A list of all of the gene groups identified by Roary [Page et al. 2015] for replicons belonging to the “pENY” plaster. Each row reports a single gene group. The “Gene” and “Annotation” values report the Prokka [Seemann 2014] predicted gene name and gene product for the genes in the group. In the case that multiple different gene names or products are predicted, all predictions are listed. The “Count” and “Pers” columns report the number and percentage of replicons in the plaster group containing members of the genome group. The remaining columns indicate the presence of the gene group in individual replicons.

**Supplemental Table S23.**

List of genes in the pAggl core genome in the context of the *Pantoea agglomerans* AR1a pAggl plasmid. Core pAggl genes were called based on prokka annotations using Roary. Where the coordinates for the annotations did not match exactly (e.g. prokka and RefSeq listed the gene with different start sites), the coordinates for RefSeq are listed as the RefSeq Annotation to be consistent with locus name, GO terms and EC numbers.

**Supplemental Table S24.**

List of genes in the pAggl core genome in the context of the *Pantoea agglomerans* AR5 pAggl plasmid. Core pAggl genes were called based on prokka annotations using Roary. Where the coordinates for the annotations did not match exactly (e.g. prokka and RefSeq listed the gene with different start sites), the coordinates for RefSeq are listed as the “RefSeq Annotation”, locus name, GO terms and EC numbers.

**Supplemental Table S25.**

A list of 125 *Pantoea agglomerans* strains was utilized in the copper tolerance assays. The presence of the *copC* gene was screened in these strains using the *copC* detection primers designed in this study. Copper tolerance was assessed based on the level of colony growth on CYE agar supplemented with varying concentrations of copper, with scores assigned as follows: 0 for no growth, 1 for non-confluent growth, and 2 for confluent growth.

**Supplemental Dataset S1.**

The consolidated and reconciled annotations from all of the WGS strains used in this study. In this study, Prokka was used, independently, by both the UGA and USDA authors to reannotate genomes for input to Roary [Page et al. 2015]. For genomes available from NCBI, both GenBank and RefSeq annotations are available. Hence, for each genome multiple sets of gene annotations are available. For each WGS strain, we have provided a table that records which locus tags in all of the available annotation sets are equivalent. That is, each row of each table, lists the locus tags from all of the genome’s available annotations sets that are equivlant. Locus tags are considered equivalent, if they refer to CDS features whose amino acid sequences are equal.

**Supplemental Document S1.**

This document details provides details of the statistical method used to analysis the leaf lesion length data.

