## Supplementary figures and images for "Plasmids encode and can mobilize onion pathogenicity in *Pantoea agglomerans*"

### Fig. S1

#### Slide 1
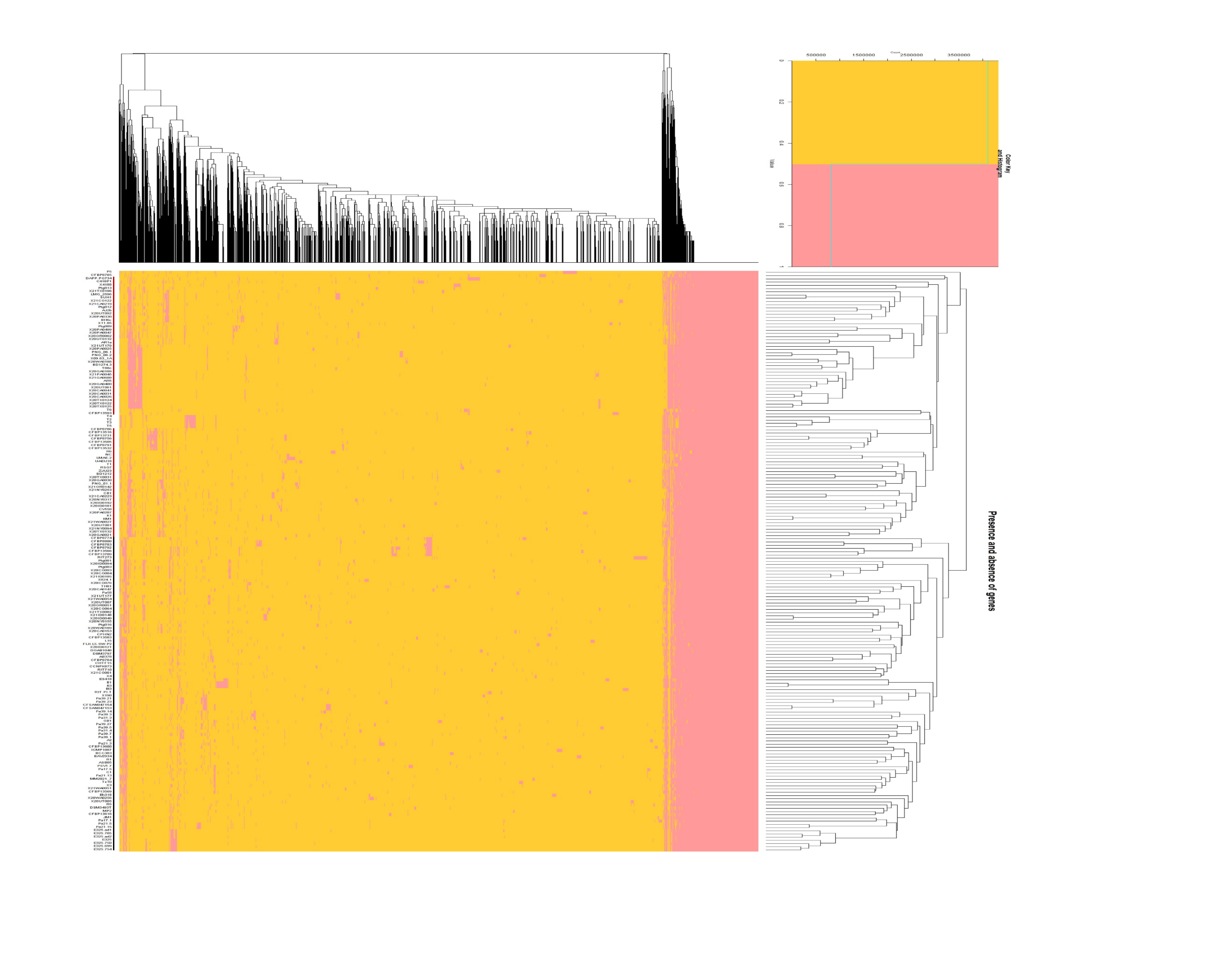

### Fig. S2

#### Slide 1
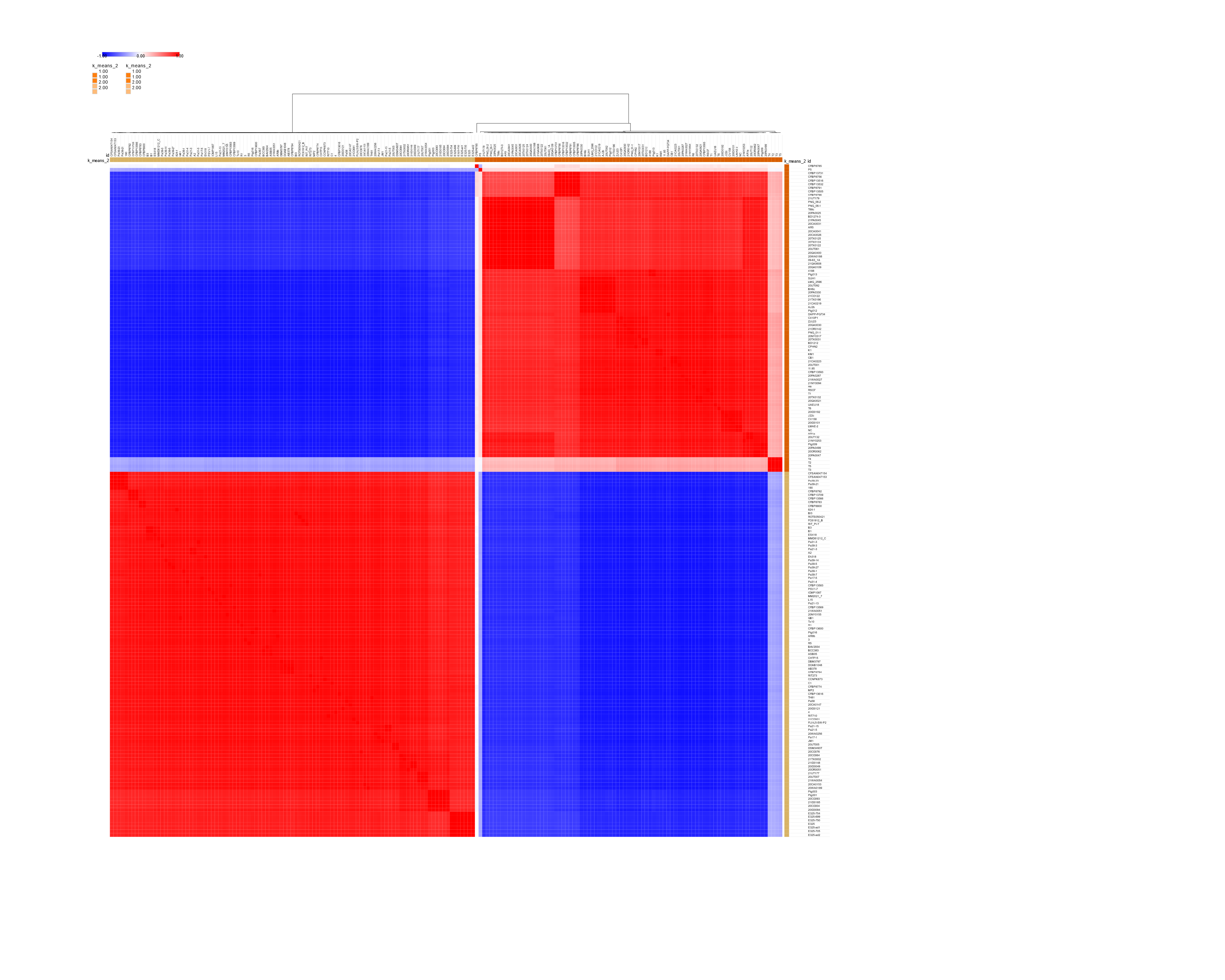

### Fig. S4

#### Slide 1
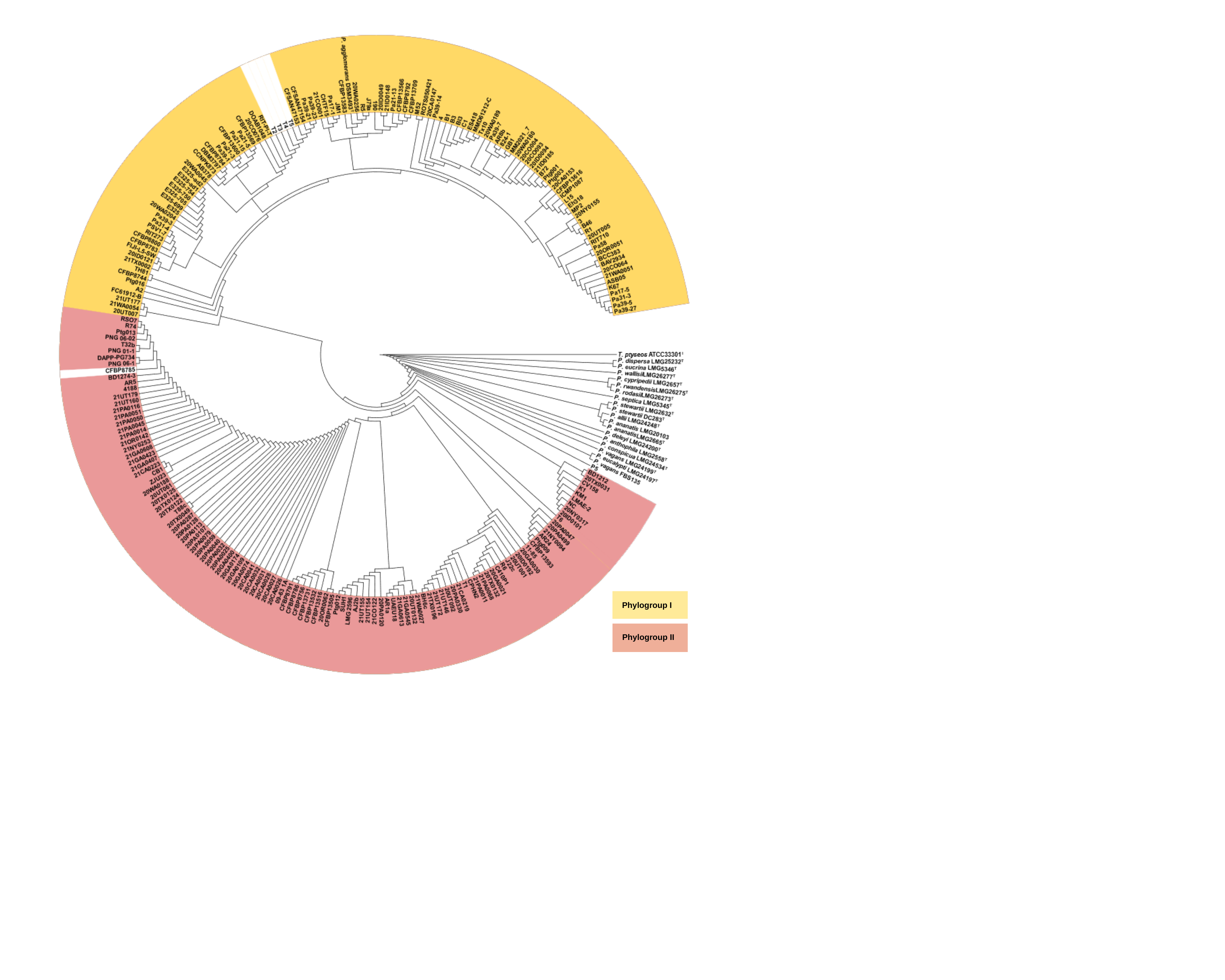

Phylogroup I
Phylogroup II

### Fig. S5

#### Slide 1
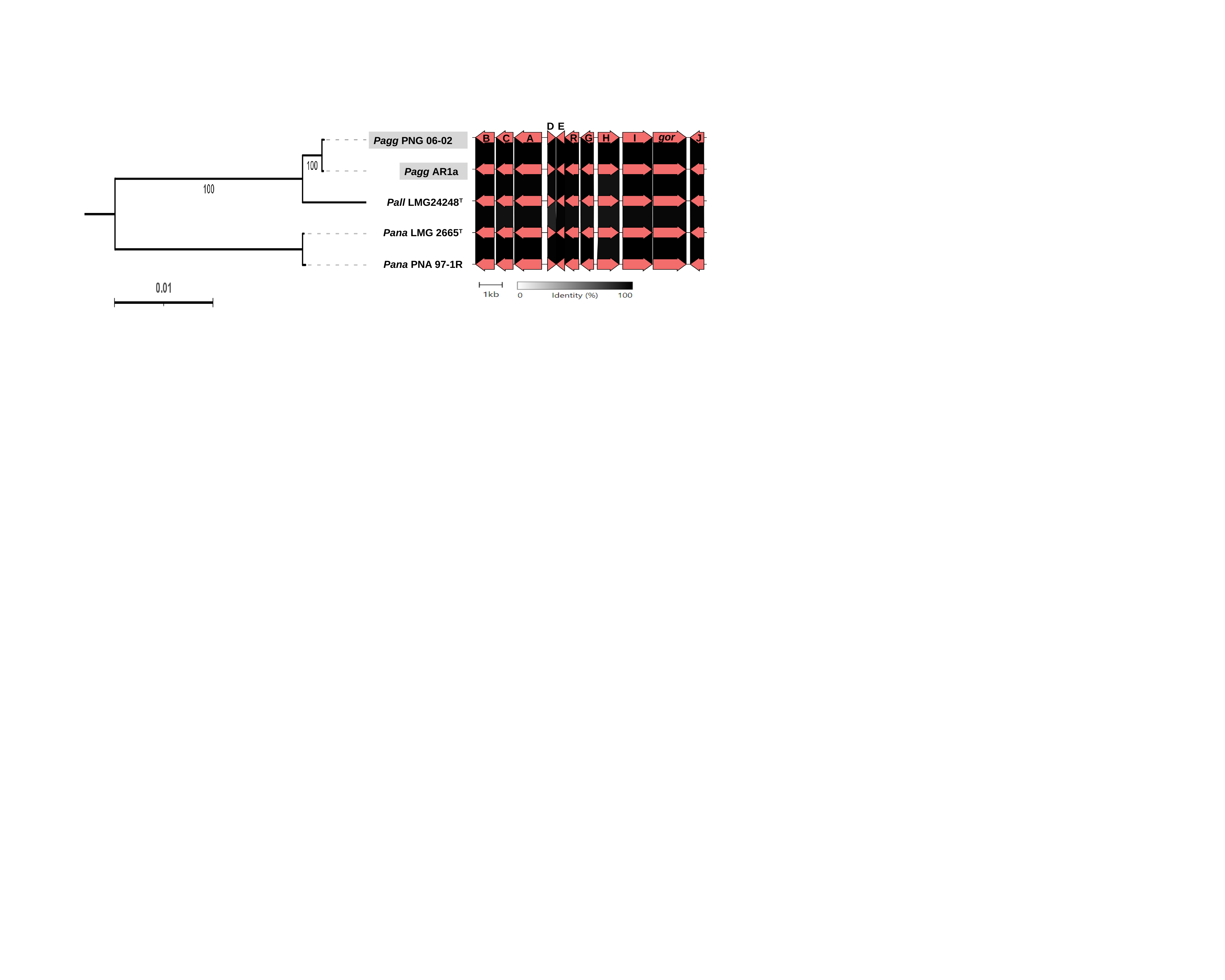

D
E
Pagg PNG 06-02
Pagg AR1a
Pall LMG24248^T^
Pana LMG 2665^T^
Pana PNA 97-1R
gor
H
J
G
I
R
A
C
B

### Fig. S6

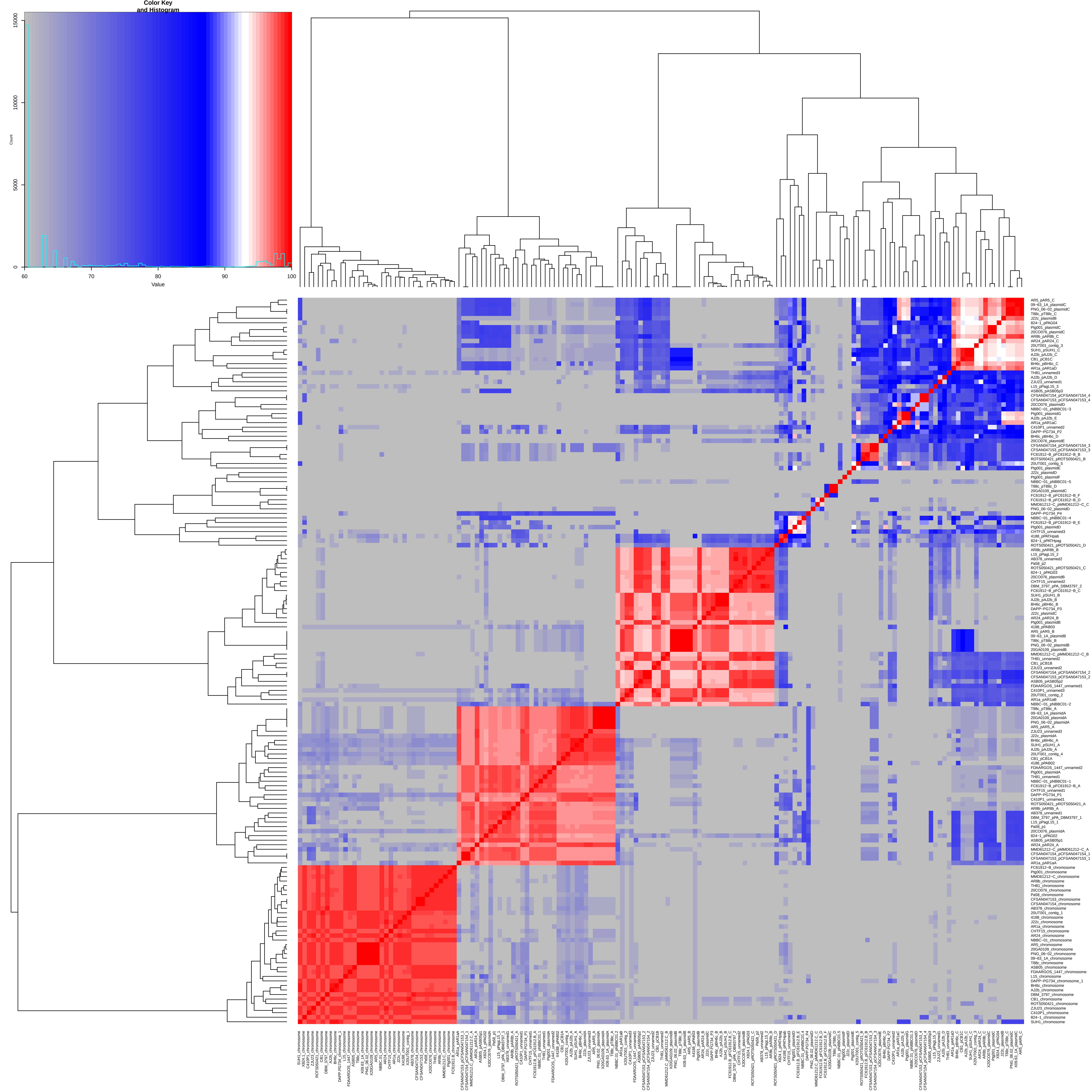

### Fig. S7

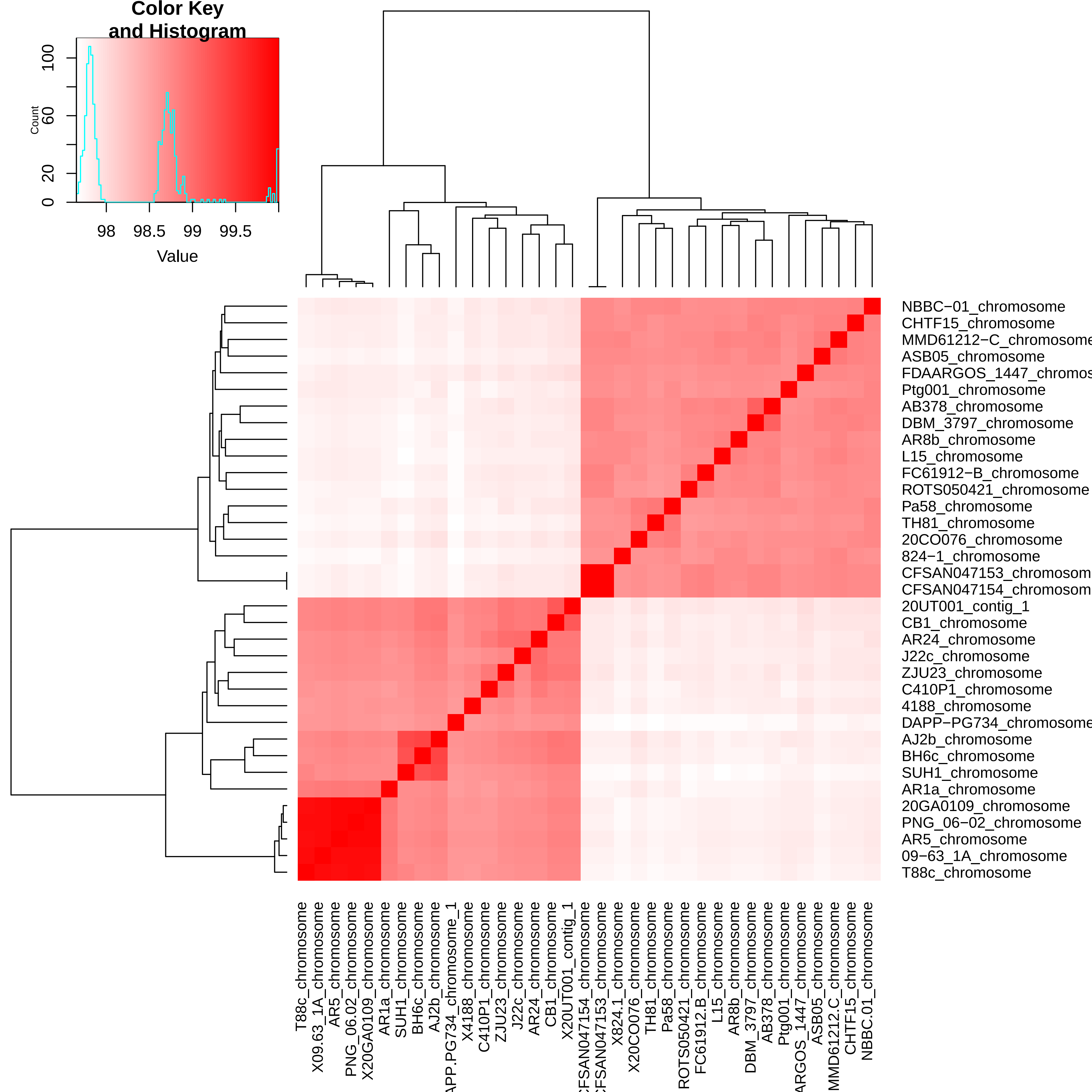

### Fig. S8

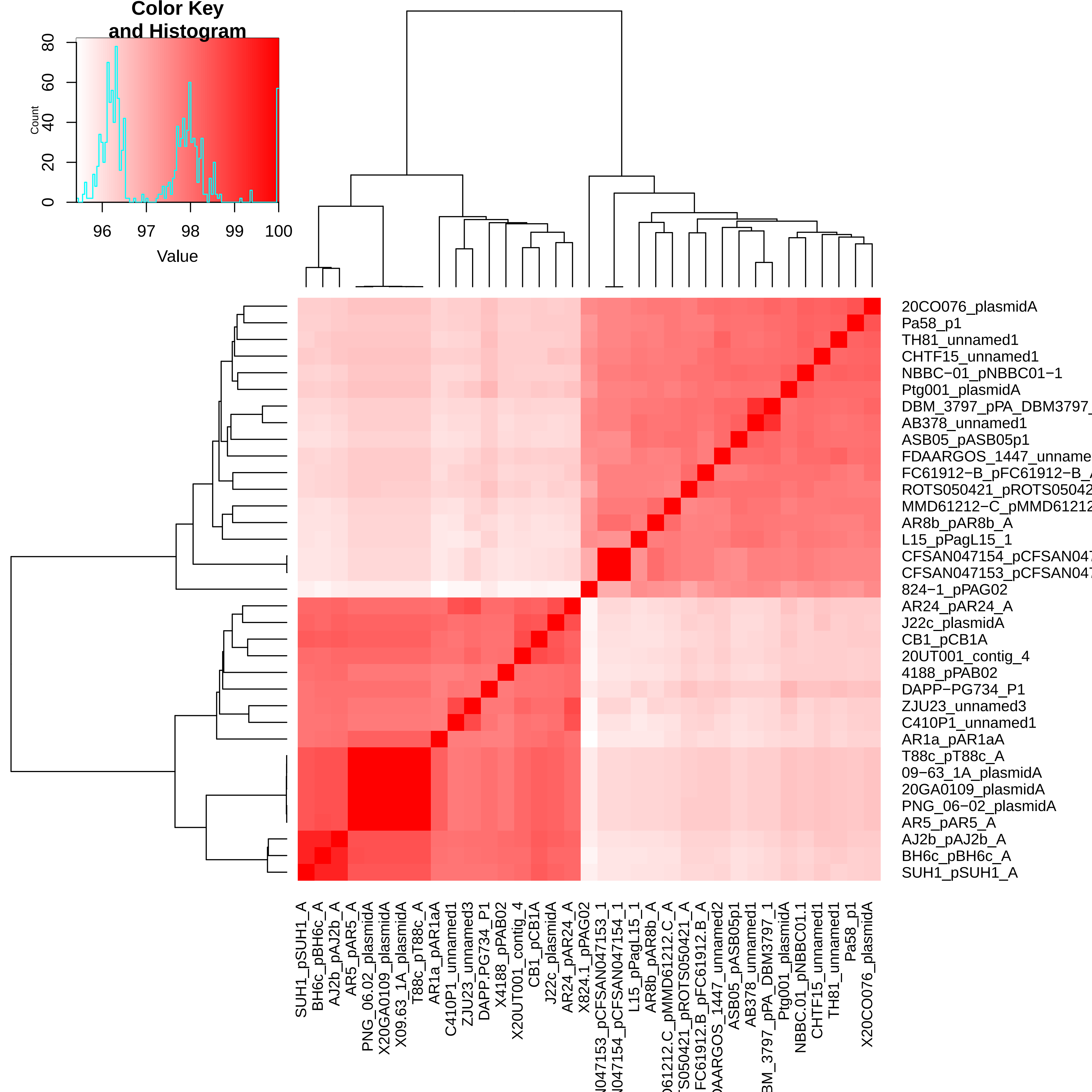

### Fig. S9

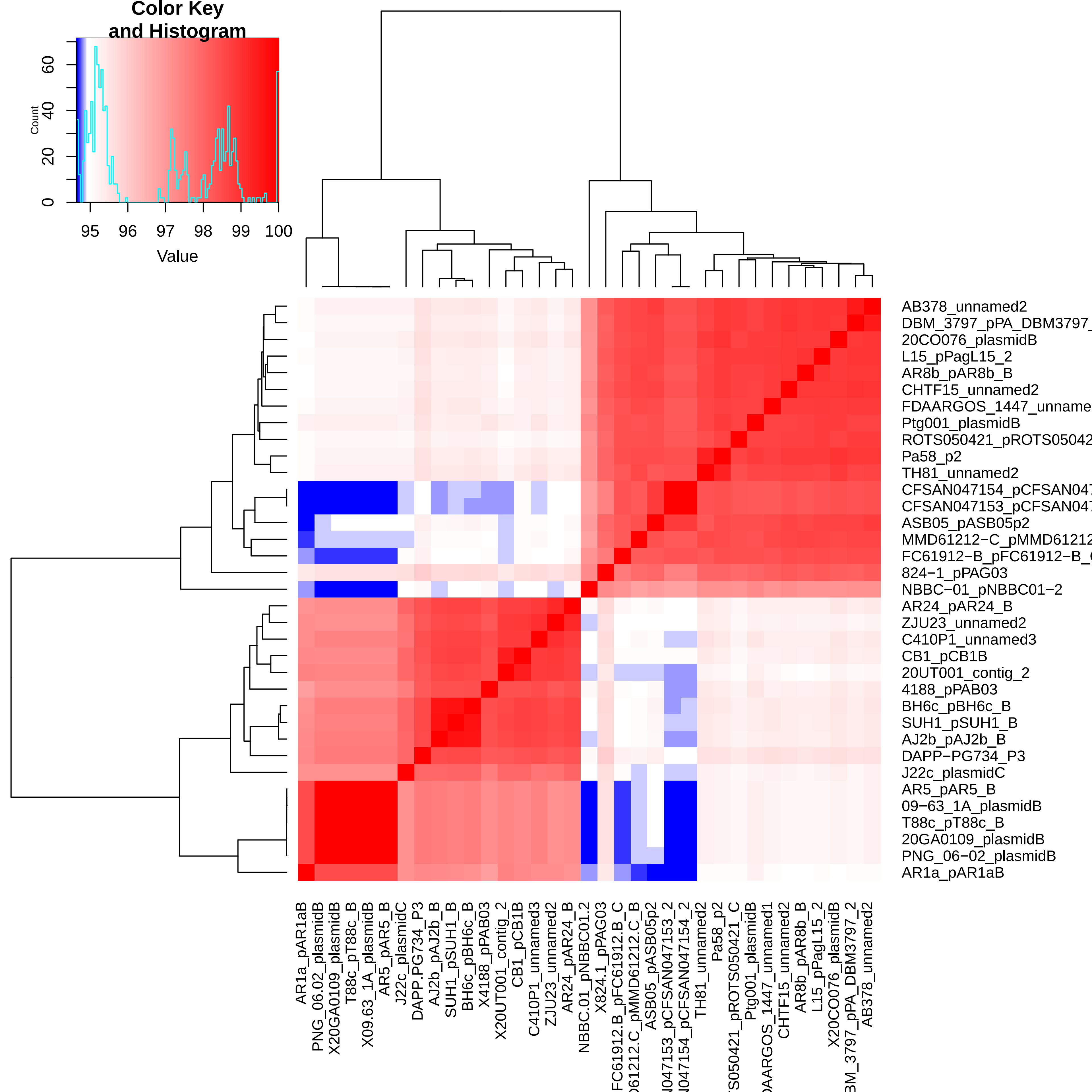

### Fig. S10

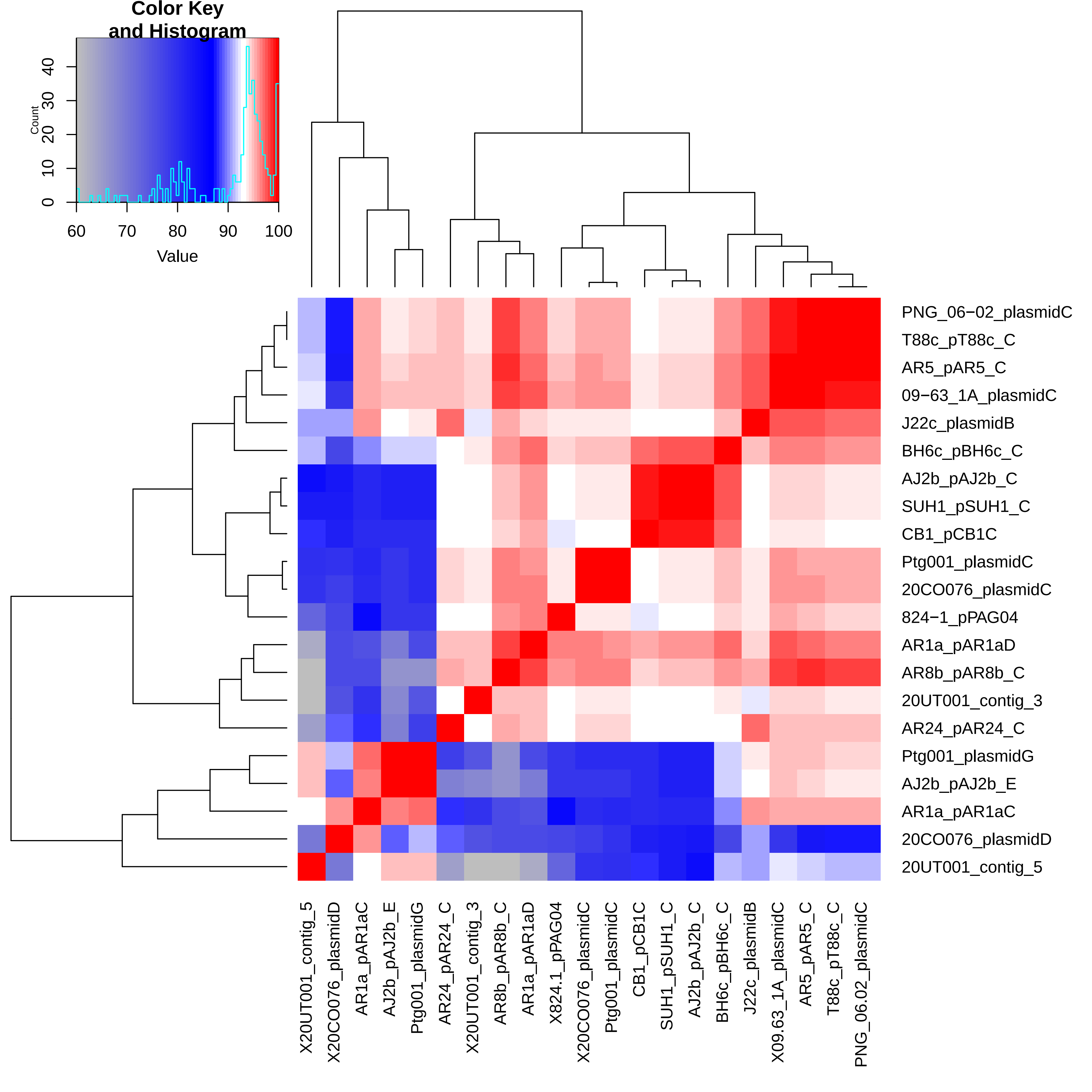

### Fig. S11

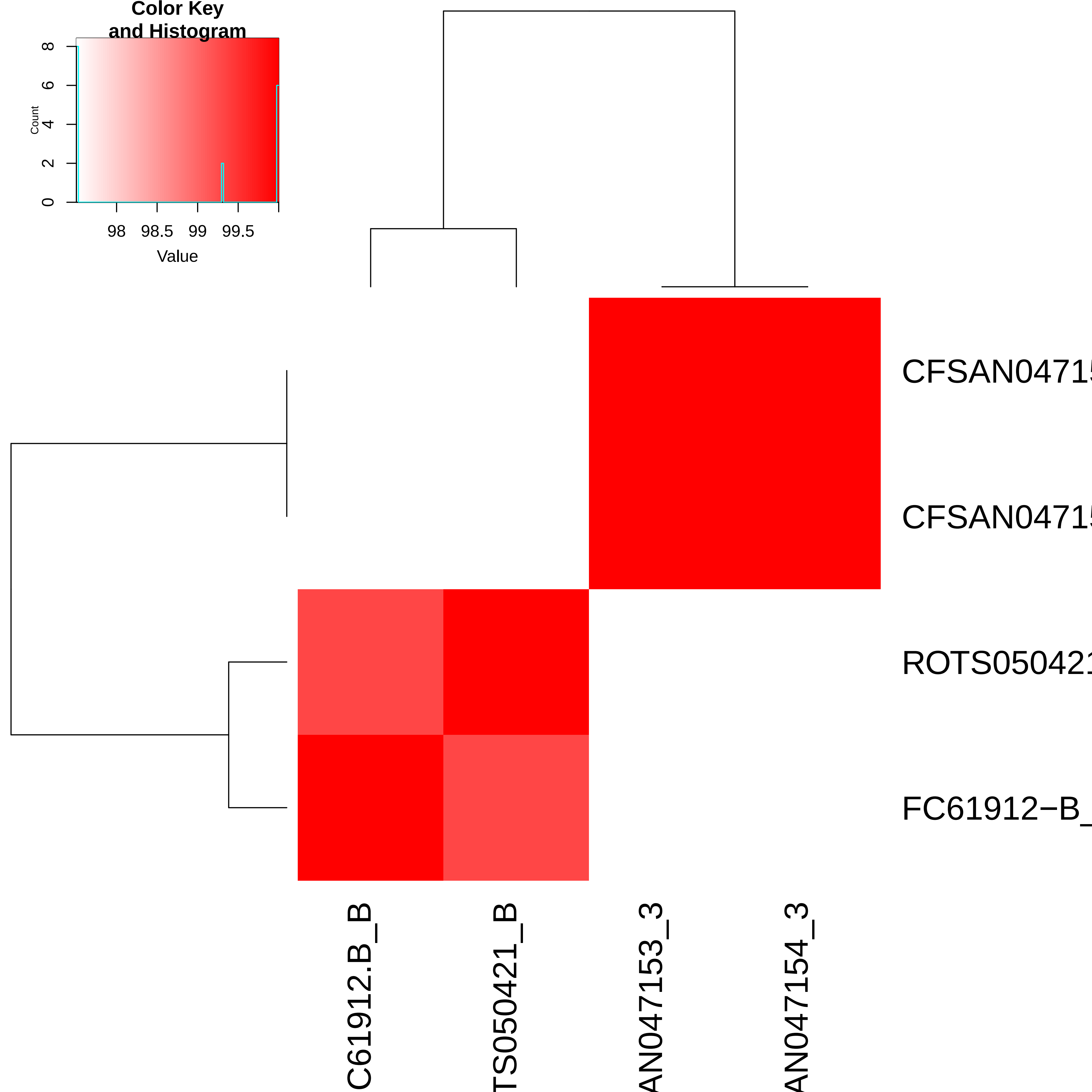

### Fig. S12

#### Slide 1
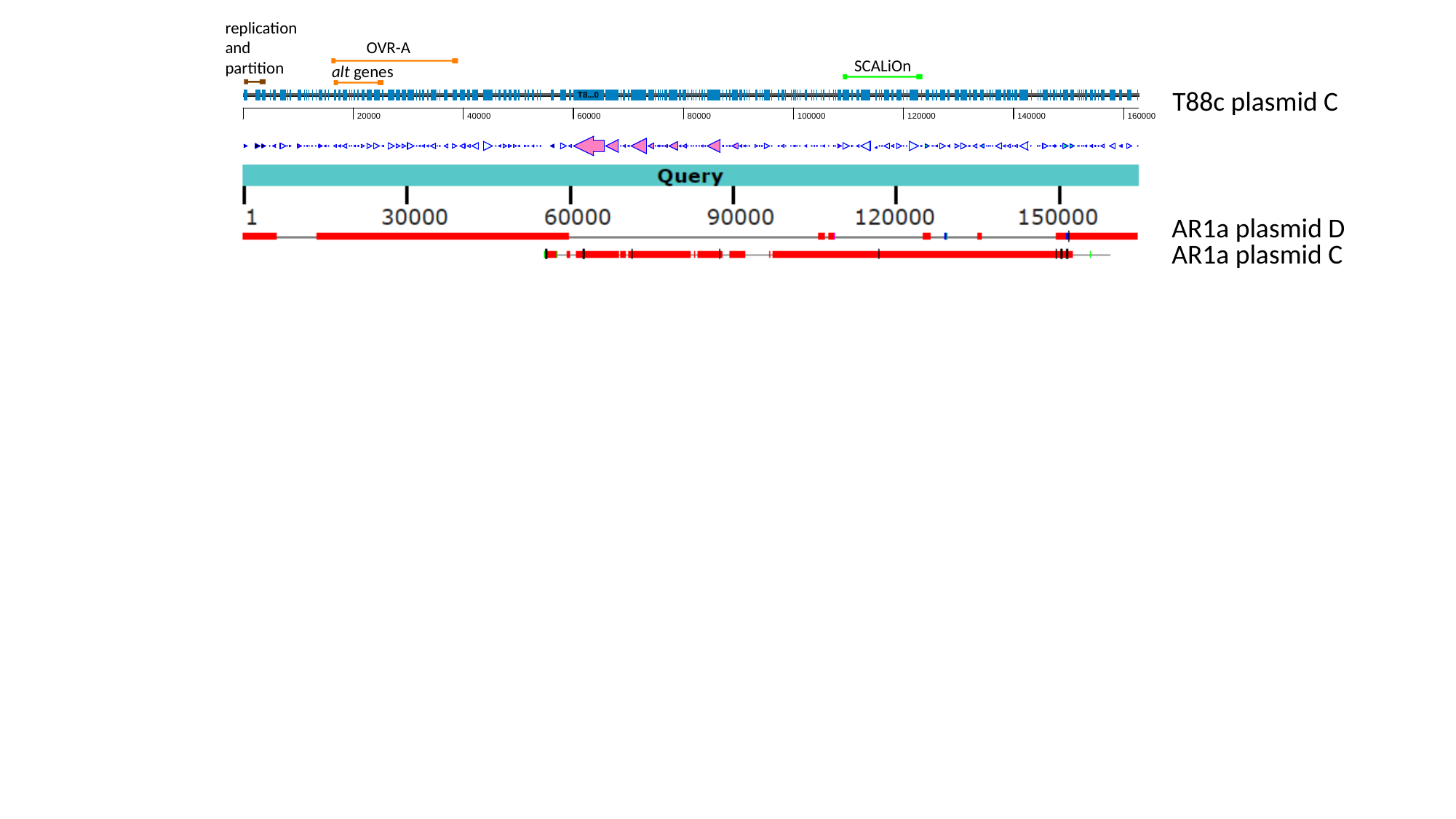

replication and partition
OVR-A
SCALiOn
alt genes
T88c plasmid C
AR1a plasmid D
AR1a plasmid C
