## Supplementary material for "Plasmids encode and can mobilize onion pathogenicity in *Pantoea agglomerans*": Fig. S3

### Slide 1
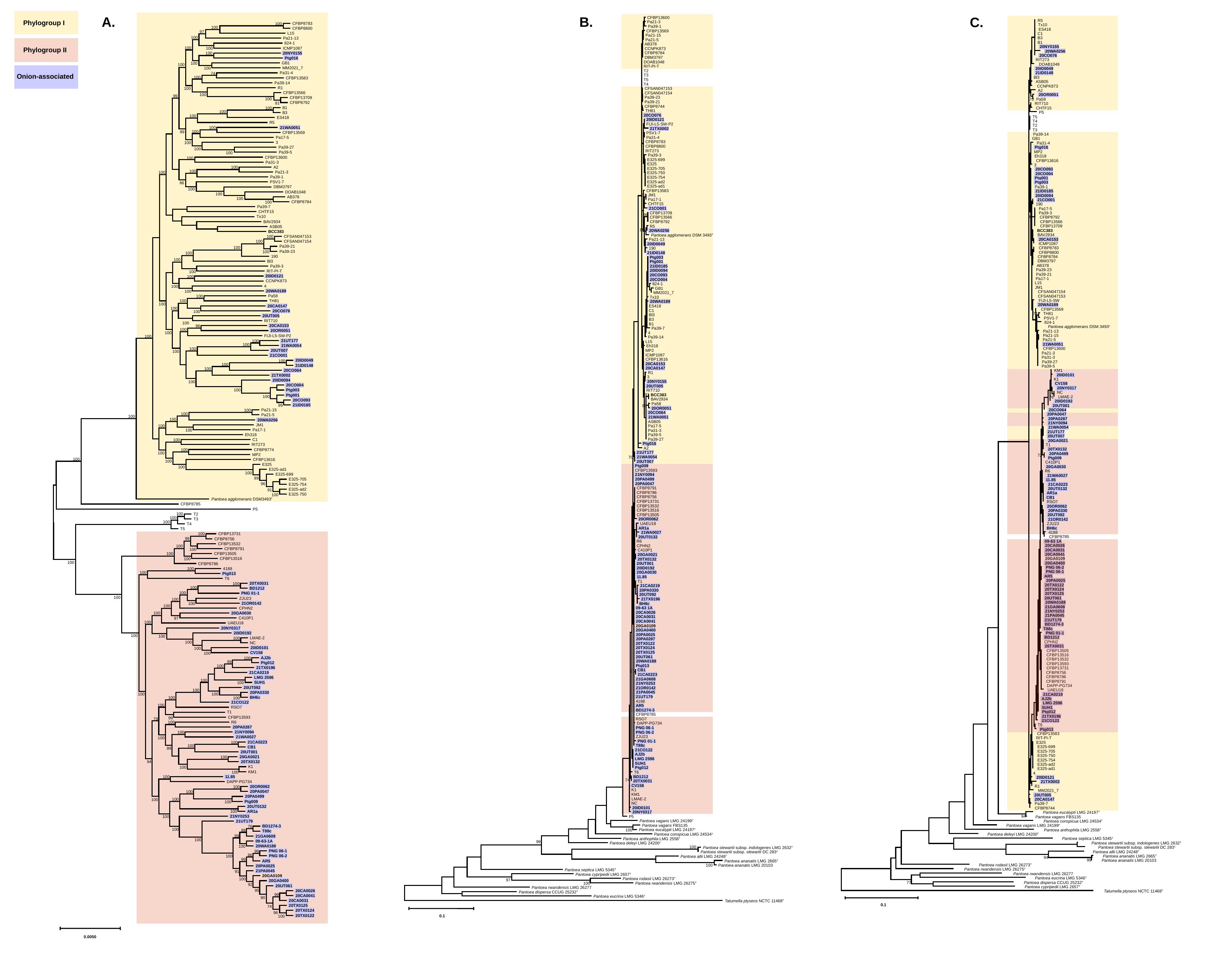

Phylogroup I
Phylogroup II
Onion-associated
A.
B.
C.
 CFBP8783
 CFBP8800
 L15
 Pa21-13
 824-1
 ICMP1087
 20NY0155
 Ptg016
 GB1
 MM2021_7
 Pa31-4
 CFBP13583
 Pa39-14
 R1
 CFBP13566
 CFBP13709
 CFBP8792
 B1
 B3
 ES418
 R5
 21WA0051
 CFBP13569
 Pa17-5
 3
 Pa39-27
 Pa39-5
 CFBP13600
 Pa31-3
 A2
 Pa21-3
 Pa39-1
 PSV1-7
 DBM3797
 DOAB1048
 AB378
 CFBP8784
 Pa39-7
 CHTF15
 Tx10
 BAV2934
 ASB05
 BCC383
 CFSAN047153
 CFSAN047154
 Pa39-21
 Pa39-23
 190
 Bl3
 Pa39-3
 RIT-PI-T
 20ID0121
 CCNPK873
 4
 20WA0189
 Pa58
 TH81
 20CA0147
 20CO076
 20UT005
 RIT710
 20CA0153
 20OR0051
 FIJI-L5-SW-P2
 21UT177
 21WA0054
 20UT007
 21CO001
100
100
97
100
100
100
100
100
100
100
74
100
100
100
99
100
81
100
100
100
100
100
99
100
100
100
100
100
100
86
100
100
100
100
100
100
100
100
100
100
96
100
100
100
100
100
100
100
100
100
100
100
100
99
 BH6c
 21CO122
 RSO7
 T1
 CFBP13593
 R6
 20PA0287
 21NY0094
 21WA0027
 21CA0223
 CB1
 20UT001
 20GA0021
 20TX0132
 K1
 KM1
 11.85
 DAPP-PG734
 20OR0062
 20PA0047
 20PA0499
 Ptg009
 20UT0132
 AR1a
 21NY0253
 21UT179
 BD1274-3
 T88c
 21GA0608
 09-63-1A
 20WA0188
 PNG 06-1
 PNG 06-2
 AR5
 20PA0025
 21PA0045
 20GA0109
 20GA0400
 20UT061
 20CA0026
 20CA0041
 20CA0031
 20TX0125
 20TX0124
 20TX0122
100
100
 20ID0049
 21ID0148
 20CO064
 21TX0002
 20ID0094
 20CO004
 Ptg003
 Ptg001
 20CO093
 21ID0185
 Pa21-15
 Pa21-5
 20WA0256
 JM1
 Pa17-1
 Eh318
 C1
 RIT273
 CFBP8774
 MP2
 CFBP13616
 E325
 E325-ad1
 E325-699
 E325-705
 E325-754
 E325-ad2
 E325-750
 Pantoea agglomerans DSM3493T
 CFBP8785
 P5
 T2
 T3
 T4
 T5
 CFBP13731
 CFBP8756
 CFBP13532
 CFBP8791
 CFBP13505
 CFBP13516
 CFBP8786
 4188
 Ptg013
 T6
 20TX0031
 BD1212
 PNG 01-1
 ZJU23
 21OR0142
 CPHN2
 20GA0030
 C410P1
 UAEU18
 20NY0317
 20ID0192
 LMAE-2
 NC
 20ID0101
 CV158
 AJ2b
 Ptg012
 21TX0196
 21CA0219
 LMG 2596
 SUH1
 20UT092
 20PA0330
100
100
100
100
100
100
100
100
100
100
100
100
100
100
100
100
100
100
100
100
100
100
100
99
96
91
100
100
100
100
100
99
100
100
100
100
100
100
100
100
100
100
100
100
100
100
100
97
100
100
100
100
100
100
100
100
99
100
100
100
100
100
100
100
100
100
99
79
100
100
100
100
100
99
100
94
100
100
100
100
100
100
100
100
100
100
76
100
88
99
100
100
99
86
100
99
91
100
92
95
99
90
74
96
100
0.0050
 CFBP13600
 Pa21-3
 Pa39-1
 CFBP13569
 Pa21-15
 Pa21-5
 AB378
 CCNPK873
 CFBP8784
 DBM3797
 DOAB1048
 RIT-PI-T
 T2
 T3
 T5
 T4
 CFSAN047153
 CFSAN047154
 Pa39-23
 Pa39-21
 CFBP8744
 TH81
 20CO076
 20ID0121
 FIJI-L5-SW-P2
 21TX0002
 PSV1-7
 Pa31-4
 CFBP8783
 CFBP8800
 RIT273
 Pa39-3
 E325-699
 E325
 E325-705
 E325-750
 E325-754
 E325-ad2
 E325-ad1
 CFBP13583
 JM1
 Pa17-1
 CHTF15
 21CO001
 CFBP13709
 CFBP13566
 CFBP8792
 R5
 20WA0256
 Pantoea agglomerans DSM 3493T
 Pa21-13
 20ID0049
 190
 21ID0148
 Ptg003
 Ptg001
 21ID0185
 20ID0094
 20CO093
 20CO004
 824-1
 GB1
 MM2021_7
 Tx10
 20WA0189
 ES418
 C1
 Bl3
 B3
 B1
 Pa39-7
 4
 Pa39-14
 L15
 Eh318
 MP2
 ICMP1087
 CFBP13616
 20CA0153
 20CA0147
 R1
 3
 20NY0155
 20UT005
 RIT710
 BCC383
 BAV2934
 Pa58
 20OR0051
 20CO064
 21WA0051
 ASB05
 Pa17-5
 Pa31-3
 Pa39-5
 Pa39-27
 Ptg016
 A2
 21UT177
 21WA0054
 20UT007
 Ptg009
 CFBP13593
 21NY0094
 20PA0499
 20PA0047
 CFBP8791
 CFBP8786
 CFBP8756
 CFBP13731
 CFBP13532
 CFBP13516
 CFBP13505
 20OR0062
 UAEU18
 AR1a
 21WA0027
 20UT0132
 R6
 CPHN2
 C410P1
 20GA0021
 20TX0132
 20UT001
 20ID0192
 20GA0030
 11.85
 T1
 21CA0219
 20PA0330
 20UT092
 21TX0196
 BH6c
 09-63 1A
 20CA0026
 20CA0031
 20CA0041
 20GA0109
 20GA0400
 20PA0025
 20PA0287
 20TX0122
 20TX0124
 20TX0125
 20UT061
 20WA0188
 Ptg013
 CB1
 21CA0223
 21GA0608
 21NY0253
 21OR0142
 21PA0045
 21UT179
 4188
 AR5
 BD1274-3
 CFBP8785
 RSO7
 DAPP-PG734
 PNG 06-1
 PNG 06-2
 ZJU23
 PNG 01-1
 T88c
87
98
85
91
80
70
87
 21CO122
 AJ2b
 LMG 2596
 SUH1
 Ptg012
 T6
 BD1212
74
 20TX0031
 CV158
 K1
 KM1
 LMAE-2
 NC
 20ID0101
 20NY0317
 P5
 Pantoea vagans LMG 24199T
 Pantoea vagans FBS135
 Pantoea eucalypti LMG 24197T
100
 Pantoea conspicua LMG 24534T
 Pantoea anthophila LMG 2558T
99
 Pantoea deleyi LMG 24200T
100
 Pantoea stewartii subsp. indologenes LMG 2632T
 Pantoea stewartii subsp. stewartii DC 283T
 Pantoea allii LMG 24248T
 Pantoea ananatis LMG 2665T
 Pantoea ananatis LMG 20103
100
 Pantoea septica LMG 5345T
 Pantoea cypripedii LMG 2657T
 Pantoea rodasii LMG 26273T
97
100
 Pantoea rwandensis LMG 26275T
 Pantoea rwandensis LMG 26277
 Pantoea dispersa CCUG 25232T
 Pantoea eucrina LMG 5346T
 Tatumella ptyseos NCTC 11468T
0.1
 R5
 Tx10
 ES418
 C1
 B3
 B1
 20NY0155
 20WA0256
 20CO076
 RIT273
 DOAB1048
 20ID0049
 21ID0148
 Bl3
 ASB05
 CCNPK873
 A2
 20OR0051
 Pa58
 RIT710
 CHTF15
 P5
 T5
 T4
 T2
 T3
 Pa39-14
 GB1
 Pa31-4
 Ptg016
 MP2
 Eh318
 CFBP13616
 3
 20CO093
 20CO004
 Ptg001
 Ptg003
 Pa39-1
 21ID0185
 20ID0094
 21CO001
 190
 Pa17-5
 Pa39-3
 CFBP8792
 CFBP13566
 CFBP13709
 BCC383
 BAV2934
 20CA0153
 ICMP1087
 CFBP8783
 CFBP8800
 CFBP8784
 DBM3797
 AB378
 Pa39-23
 Pa39-21
 Pa17-1
 L15
 JM1
 CFSAN047154
 CFSAN047153
 FIJI-L5-SW
 20WA0189
 CFBP13569
 TH81
 PSV1-7
 824-1
 Pantoea agglomerans DSM 3493T
 Pa21-13
 Pa21-15
 Pa21-5
 21WA0051
 CFBP13600
 Pa21-3
 Pa31-3
 Pa39-27
 Pa39-5
70
92
74
 KM1
 20ID0101
 K1
 CV158
 20NY0317
 NC
 LMAE-2
 20ID0192
 20UT001
 20CO064
 20PA0047
 20PA0287
 21NY0094
 21WA0054
 21UT177
 20UT007
 20GA0021
 T1
 20TX0132
 20PA0499
 Ptg009
 C410P1
 20GA0030
 R6
 21WA0027
 11.85
 21CA0223
 20UT0132
 AR1a
 CB1
 RSO7
 20OR0062
 20PA0330
 20UT092
 21OR0142
 ZJU23
 BH6c
 4188
 CFBP8785
 09-63 1A
 20CA0026
 20CA0031
 20CA0041
 20GA0109
 20GA0400
 PNG 06-2
 PNG 06-1
 AR5
 20PA0025
 20TX0122
 20TX0124
 20TX0125
 20UT061
 20WA0188
 21GA0608
 21NY0253
 21PA0045
 21UT179
 BD1274-3
 T88c
 PNG 01-1
 BD1212
 CPHN2
 20TX0031
 CFBP13505
 CFBP13516
 CFBP13532
 CFBP13593
 CFBP13731
 CFBP8756
 CFBP8786
 CFBP8791
 DAPP-PG734
 UAEU18
 21CA0219
 AJ2b
 LMG 2596
 SUH1
 Ptg012
 21TX0196
 21CO122
 T6
76
72
 Ptg013
 CFBP13583
 RIT-PI-T
 E325
 E325-699
 E325-705
 E325-750
 E325-754
 E325-ad2
 E325-ad1
 4
 20ID0121
 21TX0002
 R1
 MM2021_7
 20UT005
 20CA0147
 Pa39-7
 CFBP8744
 Pantoea eucalypti LMG 24197T
94
 Pantoea vagans FBS135
 Pantoea conspicua LMG 24534T
 Pantoea vagans LMG 24199T
 Pantoea anthophila LMG 2558T
 Pantoea deleyi LMG 24200T
 Pantoea septica LMG 5345T
Pantoea stewartii subsp. indologenes LMG 2632T
 Pantoea stewartii subsp. stewartii DC 283T
 Pantoea allii LMG 24248T
 Pantoea ananatis LMG 2665T
94
99
 Pantoea ananatis LMG 20103
 Pantoea rodasii LMG 26273T
 Pantoea rwandensis LMG 26275T
 Pantoea rwandensis LMG 26277
 Pantoea eucrina LMG 5346T
 Pantoea dispersa CCUG 25232T
77
 Pantoea cypripedii LMG 2657T
 Tatumella ptyseos NCTC 11468T
0.1
