## Supplementary material for "Plasmids encode and can mobilize onion pathogenicity in *Pantoea agglomerans*": Fig. S13

### Slide 1
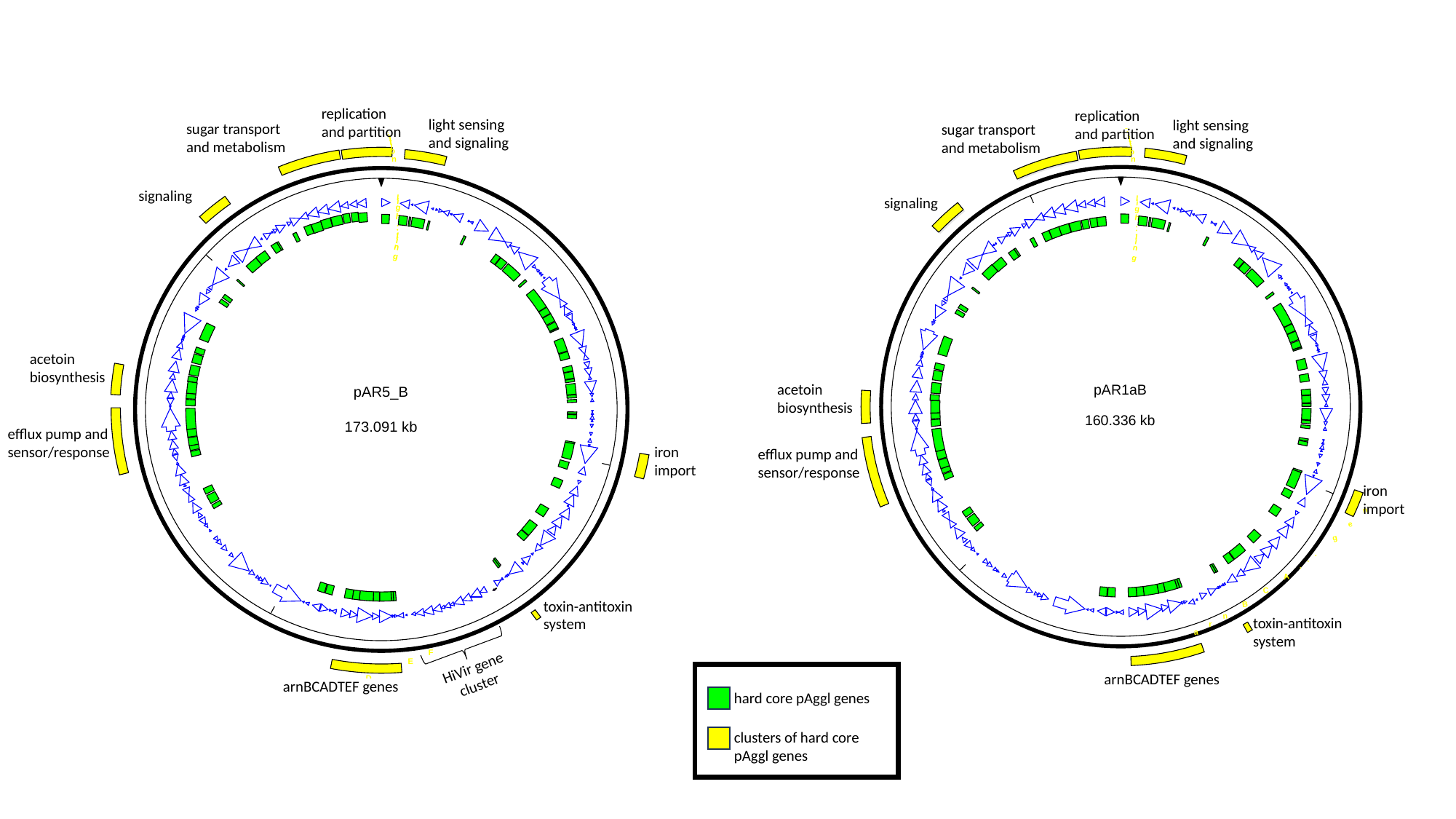

replication and partition
replication and partition
light sensing and signaling
light sensing and signaling
sugar transport and metabolism
sugar transport and metabolism
signaling
signaling
acetoin biosynthesis
acetoin biosynthesis
efflux pump and sensor/response
iron import
efflux pump and sensor/response
iron import
toxin-antitoxin system
toxin-antitoxin system
HiVir gene cluster
arnBCADTEF genes
arnBCADTEF genes
hard core pAggl genes
clusters of hard core pAggl genes
