## Supplemental Document S1 for "Plasmids encode and can mobilize onion pathogenicity in *Pantoea agglomerans*"

Description of Statistical Analysis of Leaf Assay Data

Jo Ann Asselin

For leaf assays, endpoint (5 days post inoculation) leaf lesion length was pooled from eight inoculated plants, four leaves per plant from two experiments. Lesion lengths were averaged, and standard error was plotted for error bars. In general, leaves inoculated with CB1 harboring the kanamycin-resistance cassette containing plasmid, and all of the transconjugants developed lesions over five days. None of the original strains used as recipients caused lesions in any inoculated leaves. Two of 32 water-inoculated leaves developed lesions over the course of the experiment.

Only treatments CB1 pCB1C-AKan; the transconjugants of AR8b Rp^r^, MMD61212-C Rp^r^, FC61912-B Rp^r^ (also containing pCB1C-AKan); and water were included in the model since there were no lesions (all 0 values, no variability) in other treatments. The overall effect of treatment was statistically significant.

(F_4,155_ = 57.5, p < 0.0001) Tukey post-hoc tests indicated that the mean lesion length for treatments CB1 pCB1C-AKan and the transconjugants were statistically significantly different than water, but there were no differences in CB1 pCB1C-AKan and the transconjugants at 5 days post inoculation

The 95% confidence intervals for CB1 pCB1C-AKan and the transconjugants at day 5 did not include 0, but the confidence interval for water did, indicating that water was not significantly different than 0 and not significantly different than leaves inoculated with the strains recipient strains lacking pCB1C-AKan (Figure FIG:Gina4c)
